# Herpes simplex virus 1 fluidizes the nucleus enabling condensate formation

**DOI:** 10.1101/2025.06.20.660750

**Authors:** Nora L Herzog, Tong Shu, Gururaj R Kidiyoor, Sarah Keegan, Farah Korchi, David M Chenoweth, Huaiying Zhang, Ian Mohr, Angus C Wilson, Liam J Holt

## Abstract

Molecular processes are profoundly influenced by the biophysical properties of the cell interior. However, the mechanisms that control these physical properties, and the processes they impact remain poorly understood, especially in the nucleus. We hypothesized that some viruses might change the biophysical properties of the nucleus to favor virus survival and replication and found that herpes simplex virus 1 (HSV-1) increases the mesoscale fluidity of the nucleus. The HSV-1 protein ICP4 caused fluidization and enabled growth of synthetic nuclear condensates. Conversely, conditions that decreased nuclear fluidity inhibited the growth of viral replication compartment condensates and reduced infectious virus production. Together, our data suggest that ICP4 increases nuclear fluidity to promote the formation of condensates that drive the progression of the HSV-1 life cycle. We speculate that a key function of ICP4 is to overcome the crowding and elastic confinement within cell nuclei that are a fundamental barrier to virus replication.

## INTRODUCTION

The biophysical properties of the cell interior profoundly affect biochemistry and cellular organization. Multiple factors determine cellular fluidity (the inverse of viscosity): microviscosity, macromolecular crowding, and elastic confinement decrease motion^1,2^, while non-thermal energy from metabolic and motor activities increases it^3^. The effective fluidity of the cell interior decreases in a non-linear fashion as particle size increases^4^, greatly hindering the motion of mesoscale (50-1000 nanometer length-scale) particles and cellular structures^5^. Indeed, cells are thought to border on a liquid-solid (jamming) transition at this scale, that is prevented by metabolic and motor activities driving increased motion^6,7^. These crowded, non-equilibrium properties are proposed to be tuned to enable efficient biochemistry and cellular organization: recent work shows that excessive macromolecular crowding in the cytoplasm prevents cell growth^8^ while cytoplasmic dilution leads to senescence^9^. However, despite this fundamental importance, little is known about the regulation of the physical properties of the nucleus.

Biomolecular condensate formation is crucial for biological organization^10^, and is strongly influenced by the physical environment^11^. Condensates often form through a nucleation and growth mechanism where macromolecular crowding favors nucleation but inhibits growth, while increased non-thermal energy facilitates coalescence by reducing elastic confinement and increasing the motion of condensates, thus enabling fusion^12,13^. In addition, chromatin fibers have been shown to mechanically arrest condensate growth by restricting their growth and diffusion, highlighting the importance of nuclear biophysical properties for nuclear condensate formation^14,15^.

Viruses are obligate intracellular parasites that manipulate host cells to drive viral replication. Viruses can drive extreme perturbations of cellular physiology including suppression of host cell transcription^16,17^, translation, and mRNA splicing^18^, as well as changes to chromatin architecture^19–21^ and modifications to cellular histones^22–24^. Diverse viral families form dynamic biomolecular condensates^25–27^, as part of their life cycle, including rabies virus (RABV)^28^, vesicular stomatitis virus (VSV)^29^, and severe acute respiratory syndrome coronavirus 2 (SARS-CoV-2)^30^. This led us to hypothesize that viruses might alter host cell physical properties to facilitate condensate assembly. Herpes simplex virus 1 (HSV-1) is a double-stranded DNA virus that replicates within dynamic viral condensates in the nucleus^31–36^. Therefore, we focused our studies on this virus that infects approximately two thirds of the world’s population^37^, causing oral and genital blisters, encephalitis, and neurological disability in the case of neonatal herpes^38^.

The HSV-1 life cycle consists of three phases: immediate early (IE), during which the first genes are transcribed, early (E), during which proteins required for viral DNA synthesis are produced; and late (L), which occurs after viral DNA replication has begun^39^. During the early and late phases of the HSV-1 life cycle, interchromatin domains are enlarged, enabling more efficient egress of virus capsids^40^. HSV-1 also marginalizes and condenses chromatin late in infection, in a process known as rearrangement of cellular chromatin (ROCC)^19,41^. Previous studies showed that viral replication compartments (vRCs) initially nucleate as small pre-RCs and then coalesce into larger vRC condensates^42,43^. Host DNA is excluded from vRCs during HSV-1 infection, suggesting that cellular chromatin must be manipulated to allow vRCs to form^20^. Studies examining the relationship between cellular chromatin architecture and vRC formation in HSV-1 infection correlated chromatin rearrangement with vRC expansion^41^. Taken together, this suggests that ROCC may facilitate vRC formation^44^. These studies showed that virus-directed processes including DNA synthesis, virus assembly, and virus egress may drive nuclear remodeling at the micron scale late in viral infection. However, little is known about the effect of HSV-1 on the biophysical properties of the nucleus at early times of infection, especially at the mesoscale.

Mesoscale fluidity can be studied by microrheology—the inference of the biophysical properties of materials through analysis of the motion of tracer particles. We developed nuclear Genetically Encoded Multimeric nanoparticles (nucGEMs) to enable high throughput microrheology in the nucleus^45^. HSV-1 infection increased mesoscale diffusivity at early time points of infection through a mechanism that is independent of previously described changes in nuclear architecture. Furthermore, the IE protein ICP4 was sufficient to increase mesoscale nuclear diffusivity. We showed that the ICP4-driven nuclear fluidization can facilitate condensate growth, and that inhibition of mesoscale fluidization inhibited vRC condensate maturation, leading to a decrease in infectious titer. Thus, HSV-1 fluidizes the nucleus, priming the nuclear microenvironment for optimal vRC maturation.

## RESULTS

### HSV-1 infection alters the biophysical properties of the nucleus

Until recently, microrheology has been labor intensive and slow due to difficulties in introducing tracer particles, especially to the nucleus. To overcome these limitations, we developed nuclear Genetically Encoded Multimeric nanoparticles (nucGEMs, **Fig. 1A**). nucGEMs consist of a scaffold protein fused to a fluorescent protein, which self-assembles to form a nanoparticle of defined size and shape. We made 40 nm diameter GEMs through fusion of the *Pyrococcus furiosus* encapsulin protein^46^ to mSapphire^47^. We used the SV40 nuclear localization signal to target monomers to self-assemble within the nucleus^45^. The protein monomer-encoding transgene was integrated into the genome by lentiviral transduction, leading to constitutive nucGEMexpression (**Fig. 1A**)^47^. Single particle tracking (SPT) of nucGEMs allows calculation of an effective diffusivity that reflects the effective mesoscale fluidity of the nucleoplasm at the length-scale of the particle (e.g. 40 nm). We undertook all nucGEM analysis at the 100 ms time-scale, when mesoscale effective fluidity is strongly impacted by macromolecular crowding, confinement, and non-thermal energy^6^.

**Figure 1.**
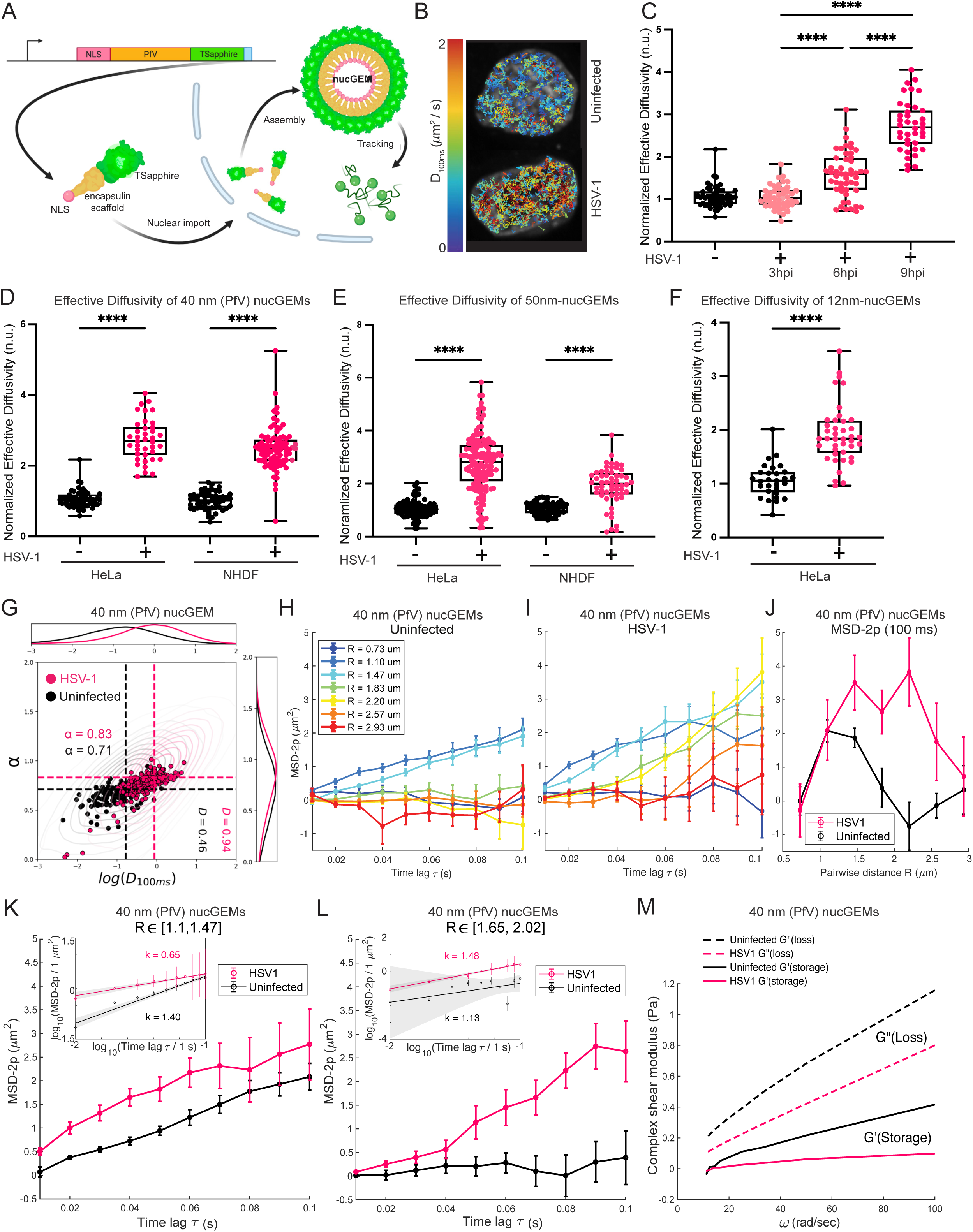
Nuclear diffusivity increases during HSV-1 infection and requires new protein synthesis. A) Schematic of the nucGEM system. (B) Individual nucGEM trajectories from representative HeLa cells +/- HSV-1, color-coded by effective diffusivity (D_eff_, see methods). (C) Effective diffusivity of nucGEMs in uninfected or HSV-1 infected HeLa cells at 3, 6, and 9 hpi. Points represent median D_eff_ of tracks in one cell, normalized to median D_eff_ of the uninfected condition. Original data were fit at 100 ms timescale, with units in µm^2^ s^-1^. See methods. n>136; N≥4 biological replicates. (D, E, F) Median D_eff_ of 40 nm GEMs (D, n>136; N≥3 biological replicates), 50nm-nucGEMs (E, n>139; N≥3 biological replicates), and 12nm-nucGEMS (F, n>69; N≥3 biological replicates) in indicated cell lines +/- HSV-1 infection at 9 hpi. (G) Scatterpolt of anomalous exponent (*α*) and log_10_ D_eff_ (*log(D_100ms_)*) of 40 nm GEM tracks in HSV-1 infected (pink) or uninfected (dark gray) cells from N≥3 biological replicates. Statistical analysis available in **Table 1**. (H, I) Two-point microrheology (TPM) of particle pairs separated by a distance range [R] in uninfected (H) and HSV-1 infected (I) NHDFs from (G). (J) TPM MSD at 100 ms as a function of R. (K, L) Direct comparison of data from (H, I) within distance ranges R [1.1 μm, 1.47 μm] (K) and R [1.65 μm, 2.02 μm] (L). Inset displays data on a log-log scale with the calculated power law exponent *k*. (M) Calculated storage modulus (G’, solid lines) and loss modulus (G”, dotted lines) in cells from (G). ****p<0.0001 by Kruskal-Wallis test (C), by two-way ANOVA (D,E) or by Mann-Whitney test (F).

To test how HSV-1 affects nucleoplasm fluidity, we expressed 40 nm diameter nuclear GEMs (nucGEMs) in HeLa cells^45^ and infected them with HSV-1 at a multiplicity of infection of 5 (MOI = 5). Cells were imaged by spinning disk confocal microscopy at 100 Hz, and SPT was performed to determine the effective diffusivity at the 100 ms timescale (D_100ms_, calculated from time-averaged mean-square displacement (MSD) **Fig. 1A-B**). Rapid acquisition limited us to 2D imaging (µm² s⁻¹). We normalized data to median control diffusivity to correct for batch effects, leading to dimensionless data (see methods). The effective diffusivity of nucGEMs increased over the course of infection, starting as early as 6 hpi (**Fig. 1C**). We next examined normal human dermal fibroblasts (NHDFs) infected with HSV-1 at MOI=5 as before and a similar increase in nucGEM diffusivity was observed in these primary cells (**Fig. 1D**).

To determine if diffusivity increased over a wider range of length scales, we additionally expressed 50 nm diameter nuclear GEMs (50nm-nucGEMs)^48^ in both HeLa cells and NHDFs and infected them as before. Effective diffusivity of 50nm-nucGEMs increased during HSV-1 infection in both cell lines (**Fig. 1E**).

To investigate physical properties at the low-end of the mesoscale, we developed 12 nm diameter nuclear GEMs (12nm-nucGEMs) based on a de novo-designed homo-60mer capsid (PDB ID: 8F53, see methods)^49^, and also observed an increase in effective diffusivity upon HSV-1 infection in HeLa cells (**Fig. 1F**). Since both NHDFs and HeLa cells showed an increase in effective diffusivity of nucGEMs, subsequent virus infection experiments were conducted in NHDFs as the more physiologically relevant cell line for HSV-1 lytic infection. Key experiments were confirmed in HeLa cells expressing 40nm-nucGEMs, in addition to cells expressing 50nm-nucGEMs and 12nm-nucGEMs to assess generality (**Fig. S1-3**).

To further characterize the nature of the changes in nucGEM motion, we graphed the log_10_ of effective diffusivity (D_100ms_) versus the anomalous exponent (**α**) (**Fig. 1G**). Shifts in **α** are indicative of changes to the type of motion—for example due to directed motion, or binding interactions with the environment—while shifts in D_100ms_ indicate changes in the effective fluidity^50,51^. The anomalous exponent and D_100ms_ both increased during HSV-1 infection, indicating increased nucGEM diffusivity and a shift toward less sub-diffusive motion. These changes are consistent with increased nucleoplasmic fluidity, likely arising from remodeling of the physical barriers that normally constrain nucGEM transport. Step size and angle distributions showed increased motion and reduced confinement after infection (**Fig. S1A-B**). Time- and ensemble-averaged MSD showed that **α** increased (from 0.85 to 0.94) while D doubled from 0.61 to 1.2 µm^2^ s^-1^ in control versus infected conditions, further supporting the conclusion that nucleoplasmic fluidity increases during HSV-1 infection concomitantly with nuclear remodeling (**Fig. S1C**). Distributions of log_10_ of effective diffusivity versus **α** during HSV-1 infection with 50nm-nucGEMs (**Fig. S1D**) and 12nm-nucGEMs (**Fig. S1E**) showed similar effects to the 40nm-nucGEM data. Thus, HSV-1 infection leads to an increase in nucleoplasmic fluidity, likely by reducing barriers to particle motion across multiple length-scales.

To more deeply characterize how biophysical properties of the nucleus changed during infection, we employed two-point microrheology (TPM). TPM analyzes correlations in pairwise particle movement and detects coupling of motion, for example due to flows^52,53^. Motion was correlated only at distances greater than 0.73 µm and less than 1.83 µm in uninfected NHDF cells (**Fig. 1H**). HSV-1 infection led to correlations at longer distances, up to 2.57 µm (**Fig. 1I**); **Fig. 1J** shows a pairwise comparison at 100 ms time lag. Focusing on two fixed distance ranges, we see that at 1.10 - 1.47 µm correlated motion drives clear increases in 2-point MSD for both infected and uninfected cells (**Fig. 1K**), while at 1.65 - 2.02 µm separation, 2-point MSD still increases for infected cells, but uncorrelated motion leads to a flat 2-point MSD near zero in uninfected cells (**Fig. 1L**). At these larger distances, HSV-1 infection also increased the power-law exponent (*k*) to values greater than 1, indicating that active intracellular processes drive correlated motion at this scale^53,54^.

Finally, to further examine elastic confinement, we assessed the velocity autocorrelation of nucGEM trajectories under infected and uninfected conditions (**Fig. S1F-G**) and saw that all trajectories showed a negative velocity autocorrelation at short timescales, consistent with elastic behavior in the nucleoplasm^55^. The value of this autocorrelation was variable dependent on timescale (**Fig. S1G, inset**), also consistent with elastic behavior. Using the generalized Stokes-Einstein relation for complex viscoelastic materials^56,57^, we evaluated the frequency-dependent rheological properties of the nucleoplasm to infer the storage modulus (G’, characterizing elastic behavior) and loss modulus (G”, characterizing viscous behavior) in uninfected and HSV-1 infected cells (**Fig. 1M**). Our data showed the loss modulus was greater than the storage modulus under both conditions, indicating the nucleoplasm behaves as a viscoelastic fluid under both infected and uninfected conditions^56^. Both the storage modulus and loss modulus were reduced during HSV-1 infection compared to control conditions (**Fig. 1M**), indicating that both elasticity and viscosity were reduced in the nucleoplasm during HSV-1 infection, consistent with fluidization.

### HSV-1 transcription and translation are required to increase nucGEM diffusivity

To determine if HSV-1 tegument or gene expression was required to increase nucGEM diffusivity, we infected cells with UV-inactivated HSV-1. The tegument (a protein layer between envelope and capsid^58^) remains intact during UV-inactivation, which prevents viral transcription but allows capsid entry. UV-inactivated HSV-1 neither expressed viral gene products nor increased nuclear diffusivity (**Fig. S1H-J**). Similarly, cycloheximide treatment, which permits IE RNA accumulation but blocks protein synthesis, prevented increased nuclear diffusivity (**Fig. S1K-M**).

### Fluidization is independent of viral DNA synthesis, chromatin margination, and nuclear volume changes

HSV-1 DNA replication is linked to architectural rearrangement of the infected nucleus, most notably the condensation and margination of cellular chromatin^19,20,41,44^. Previous studies linked capsid diffusivity changes to these large-scale rearrangements^40,41,59^. Therefore, we asked if virus DNA replication was required to increase nucGEM diffusivity. We blocked virus DNA replication either genetically, using a mutant virus deficient for the essential single-stranded DNA binding protein ICP8 (d301, henceforth referred to as ΔICP8)^60^, or by inhibiting the viral DNA polymerase with phosphonoacetic acid (PAA)^61^. Unexpectedly, nucGEM diffusivity increased even when virus DNA replication was prevented or inhibited (**Fig. 2A-B, Fig. S2A,C**). Without DNA replication (ΔICP8 or PAA), chromatin margination was absent, indicating mesoscale fluidization occurs independently of large-scale rearrangements (**Fig. 2C**). Complex shear modulus analysis^56,57^ confirmed ΔICP8 infection decreased nucleoplasm elasticity and viscosity, similar to WT HSV-1 (**Fig. S2D**).

**Figure 2.**
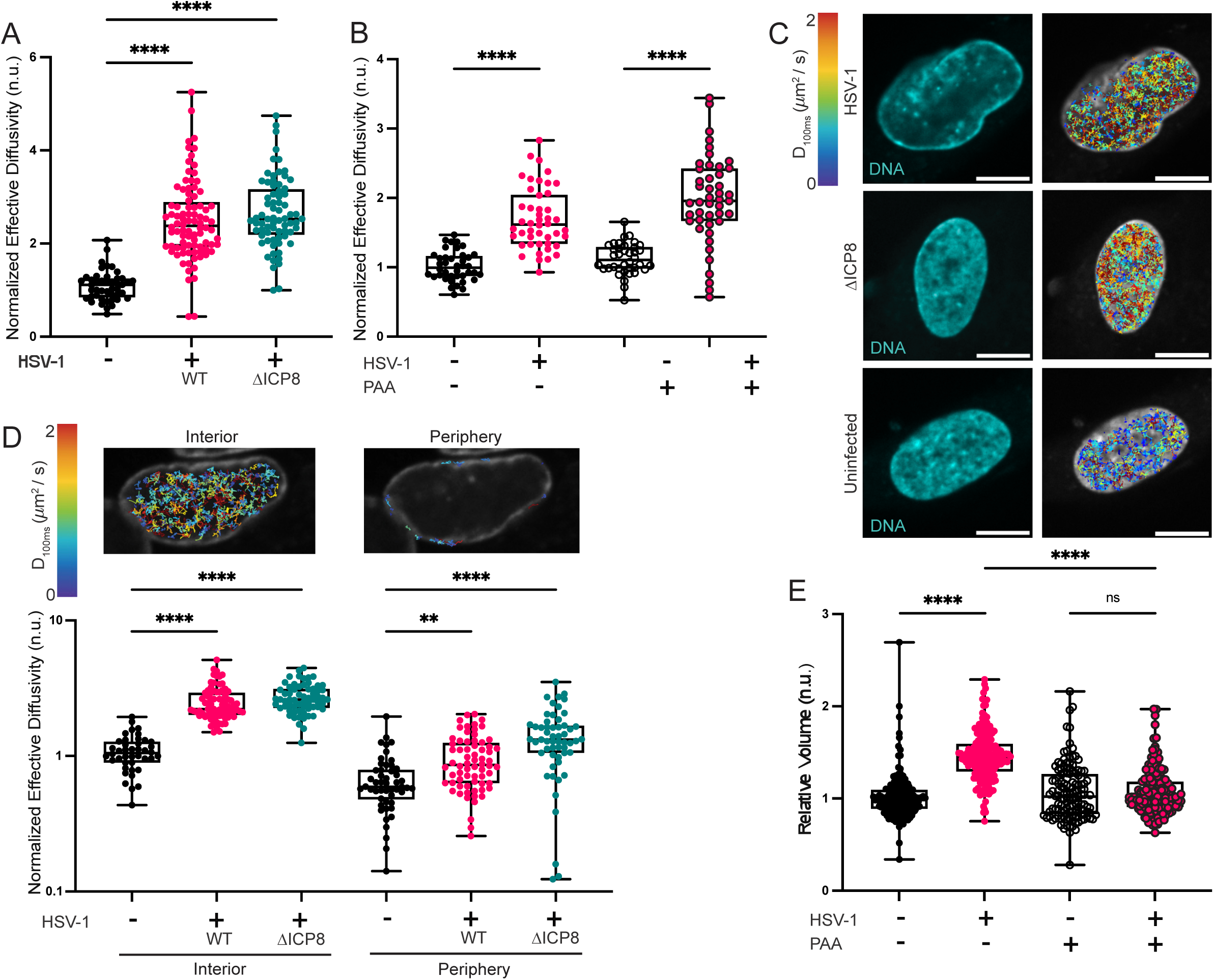
HSV-1 infection fluidizes the nucleus independent of viral DNA synthesis, chromatin margination, and nuclear volume changes. (A) Effective diffusivity of nucGEMs in NHDFs infected with WT HSV-1 or ΔICP8 at 9 hpi, MOI = 5. n>83; N≥3 biological replicates. (B) Effective diffusivity of NHDFs 9 hpi with WT HSV-1, treated with 300 μg/mL phosphonoacetic acid (PAA) or untreated. n>86; N≥3 biological replicates. (C) Cells from (A) were stained with SiR-DNA, which binds to AT-rich DNA, to visualize the nucleus (left). Individual nucGEM trajectories, color-coded by effective diffusivity, were projected onto the nuclear images (right). Scale bar, 10 μm. (D) nucGEM diffusivity in the periphery and the interior of cells from (A). Periphery is the outer 1830 nm of the nucleus. n>80; N≥3 biological replicates. (E) Relative nuclear volume of cells from (B). All volumes normalized to the median uninfected volume by experiment. n>249; N≥2 biological replicates. IF images are representative of N≥3 biological replicates. **p<0.01, ****p<0.0001 by Kruskal-Wallis test (A) or by two-way ANOVA (B,D,E).

We next explored spatial regulation of nuclear properties. Chromatin in the periphery of the nucleus tends to be more compact than that in the center^59^. Therefore, we tested whether nucGEM motion was distinct at the periphery compared to the interior of the nucleus. We compared nucGEM motion in an outer ring of 1800 nm width at the periphery of the nucleus (the approximate extent of marginated chromatin in NHDFs during WT HSV-1 infection) to the region interior to this ring. Diffusivity was lower in the periphery than the interior for all conditions, but both compartments were fluidized to a similar extent during WT and ΔICP8 infection (**Fig. 2D**).

Finally, we measured nuclear volume, which increases during HSV-1 infection and could reduce crowding^62^. Nuclear volume increased during WT infection but not during ΔICP8 infection or PAA treatment (**Fig. 2E**), indicating that nuclear fluidization during early HSV-1 infection does not require increased nuclear volume.

### The HSV-1 immediate early protein ICP4 is required to fluidize the infected cell nucleoplasm

Since late genes require DNA replication^58^ but fluidization occurs without it, we reasoned an IE or early protein drives increased fluidity. We infected NHDFs with viruses lacking each nuclear IE protein (ICP0, ICP4, ICP22, or ICP27) and measured nucGEM diffusivity. ΔICP0, ΔICP22, and ΔICP27 viruses increased diffusivity by 9 hpi, but ΔICP4 did not (**Fig. 3A**). Viruses lacking ICP0, ICP22, or ICP27 all expressed ICP4 as measured by immunoblot (**Fig. S3A**). Thus, nuclear fluidization requires ICP4.

**Figure 3.**
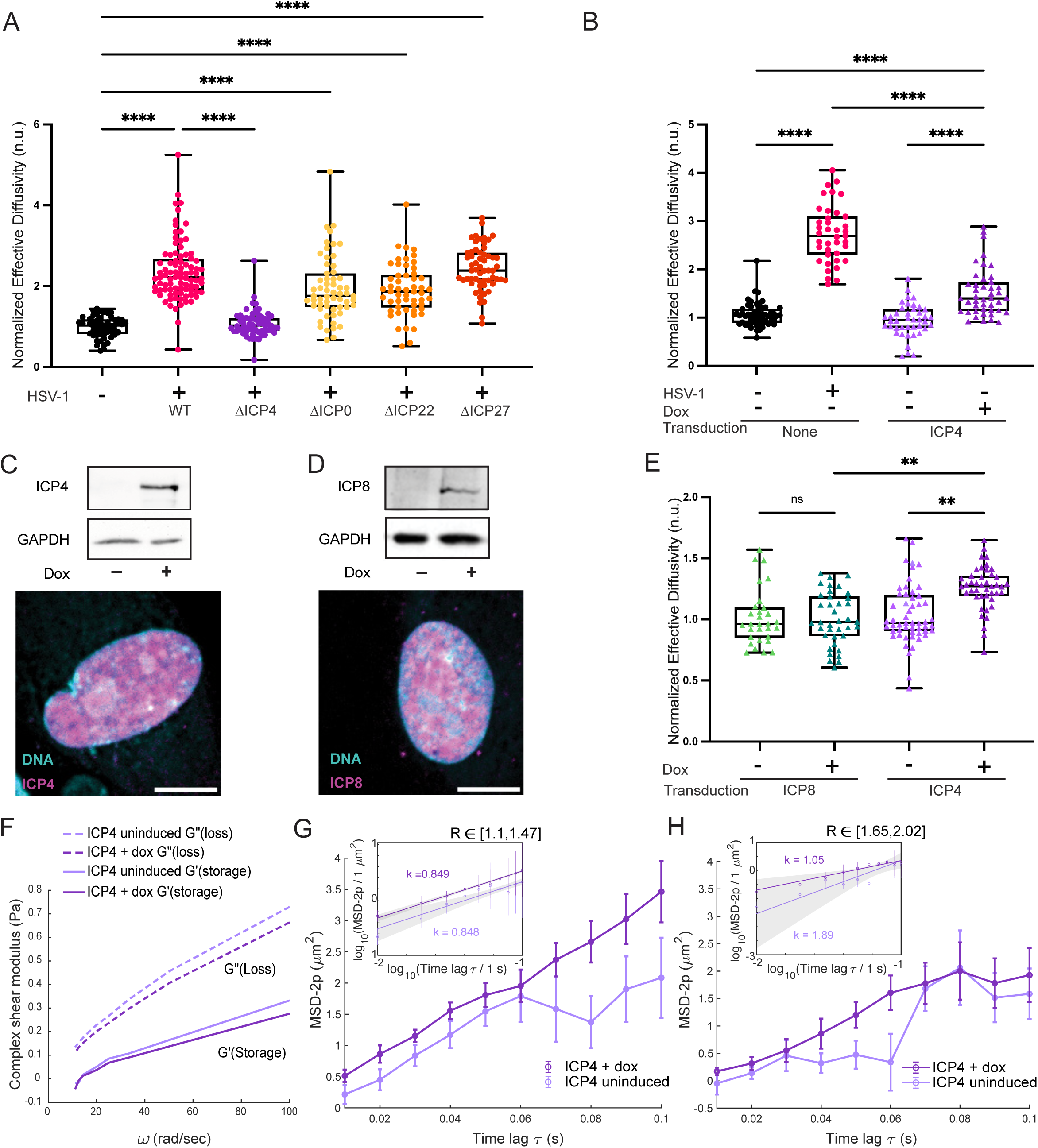
HSV-1 immediate early protein ICP4 is necessary to fluidize the infected cell nucleoplasm and sufficient to fluidize the nucleoplasm of uninfected cells. (A) Effective diffusivity (D_eff_) of nucGEMs in NHDFs infected with WT HSV-1, ΔICP4, ΔICP22, or ΔICP27 at an MOI of 5, or with ΔICP0 at an MOI of 10, at 9 hpi. n>108; N≥3 biological replicates. (B) D_eff_ of nucGEMs in HeLa cells infected with WT HSV-1 or with exogenous ICP4 induction for 9 h. n>81; N≥4 biological replicates. (C) Immunoblot and IF for ICP4 or (D) ICP8 induction in NHDFs at 9 hpt. Scale bar, 10 μm. (E) D_eff_ of nucGEMs in NHDFs with exogenous expression of ICP4 or ICP8 for 9 h. n>107; N≥5 biological replicates. (F) Calculated storage modulus (G’, solid lines) and loss modulus (G”, dotted lines) in cells identified in (E). (G, H) TPM from cells identified in (E) within distance ranges R [1.1 μm, 1.47 μm] (G) and R [1.65 μm, 2.02 μm] (H). Inset displays data on a log-log scale with the calculated power law exponent *k*. IF images are representative of N≥3 biological replicates. **p<0.01, ****p<0.0001 by Kruskal-Wallis test.

We further confirmed that fluidization was independent of viral DNA replication, but dependent on ICP4 by graphing the ensemble-time-averaged MSD as previously, and showed an increase in **α** and D in both during infection by WT HSV-1 and ΔICP8 (**Fig. S3B**), but not ΔICP4. The velocity autocorrelation value at **τ** = 0.01 s also decreased during infection with WT HSV-1 or ΔICP8 (**Fig. S3C**) compared to uninfected cells but these differences were not statistically significant. Moreover, there was no change in velocity autocorrelation compared to control during infection with ΔICP4 virus. These results are consistent with a model where ICP4 is required for mesoscale fluidization of the nucleoplasm during HSV-1 infection.

### ICP4 is sufficient to fluidize the nucleus of uninfected cells

To test whether ICP4 alone suffices for increased diffusivity, we transfected HeLa cells with doxycycline-inducible ICP4. Exogenous expression of ICP4 led to significantly lower levels of protein expression than during HSV-1 infection (**Fig. 3B, Fig. S3D**). Nevertheless, even this low level of exogenous ICP4 was sufficient to increase nucGEM diffusivity in HeLa cells after 9h of induction. For NHDFs, we constructed lentiviral vectors expressing inducible ICP4 or control protein ICP8. The efficacy of ICP4 and ICP8 expression and their appropriate nuclear localization were verified by immunoblot and indirect immunofluorescence (IF) respectively (**Fig. 3C-D**). ICP4 induction increased nucGEM diffusivity in primary NHDFs, while ICP8 induction had no detectable effect on nucGEM diffusivity (**Fig. 3E**). Furthermore, induction with ICP4 was sufficient to decrease both the elasticity and viscosity of the nucleoplasm, again consistent with nuclear fluidization (**Fig. 3F**). The ability of exogenous ICP4 expression to increase nucleoplasm fluidity at multiple length-scales was confirmed using 50nm-nucGEMs and 12nm-nucGEMs (**Fig. S3E-F**). Thus, exogenous expression of ICP4, but not another HSV-1 nuclear protein ICP8, is sufficient to partially recapitulate the nuclear fluidization occurring during HSV-1 lytic infection.

To confirm the biophysical fluidization of the nucleus by exogenous ICP4 expression, we again examined the angle distribution (**Fig. S3G**) and the log_10_ of effective diffusivity versus **α** (**Fig. S3H-I**), which both show shifts in the same direction during exogenous ICP4 induction as seen during HSV-1 infection, indicating ICP4 induction is altering nucleoplasm fluidity and particle confinement. Both storage and loss moduli of the nucleoplasm were decreased upon ICP4 expression compared to control conditions (**Fig. 3F**), and velocity autocorrelation indicated a small decrease in elastic confinement upon ICP4 induction (**Fig. S3K-M**). Finally, using TPM, we verified that ICP4 induction leads to a higher 2-point MSD than in uninduced cells at multiple particle distances (**Fig. 3G-H**). Uninduced cells appeared to behave similarly to uninfected cells at short timescales, but showed higher 2-point MSDs at longer timescales, similar to HSV-1 infection.

### Both the N-terminal and C-terminal domains of ICP4 are required to increase nucGEM diffusivity in uninfected cells

We used mutants to identify ICP4 domains required for fluidization: N-terminal activation domain (NTA), C-terminal activation domain (CTA), and DNA-binding domain (DBD)^63^. We engineered three mutants (**Fig. 4A**): CTA deletion (ICP4-ΔCTA)^64^, NTA truncation (ICP4-ΔNTA)^64^, and DBD mutation (ICP4-mDBD, R456L/R457L)^65^. This mutation in the DBD prevents specific binding of a sequence found in ICP4-regulated promoters, but does not prevent non-specific binding of ICP4 to host chromatin^63,65,66^. All mutants expressed at comparable levels to WT ICP4 and localized to nuclei (**Fig. 4B-C**). After 9h induction in nucGEM-expressing NHDFs, ICP4-mDBD increased diffusivity like WT ICP4, but ICP4-ΔNTA and ICP4-ΔCTA did not (**Fig. 4D**). We did not detect changes in nuclear volume between induced and uninduced cells (**Fig. S4A**).

**Figure 4.**
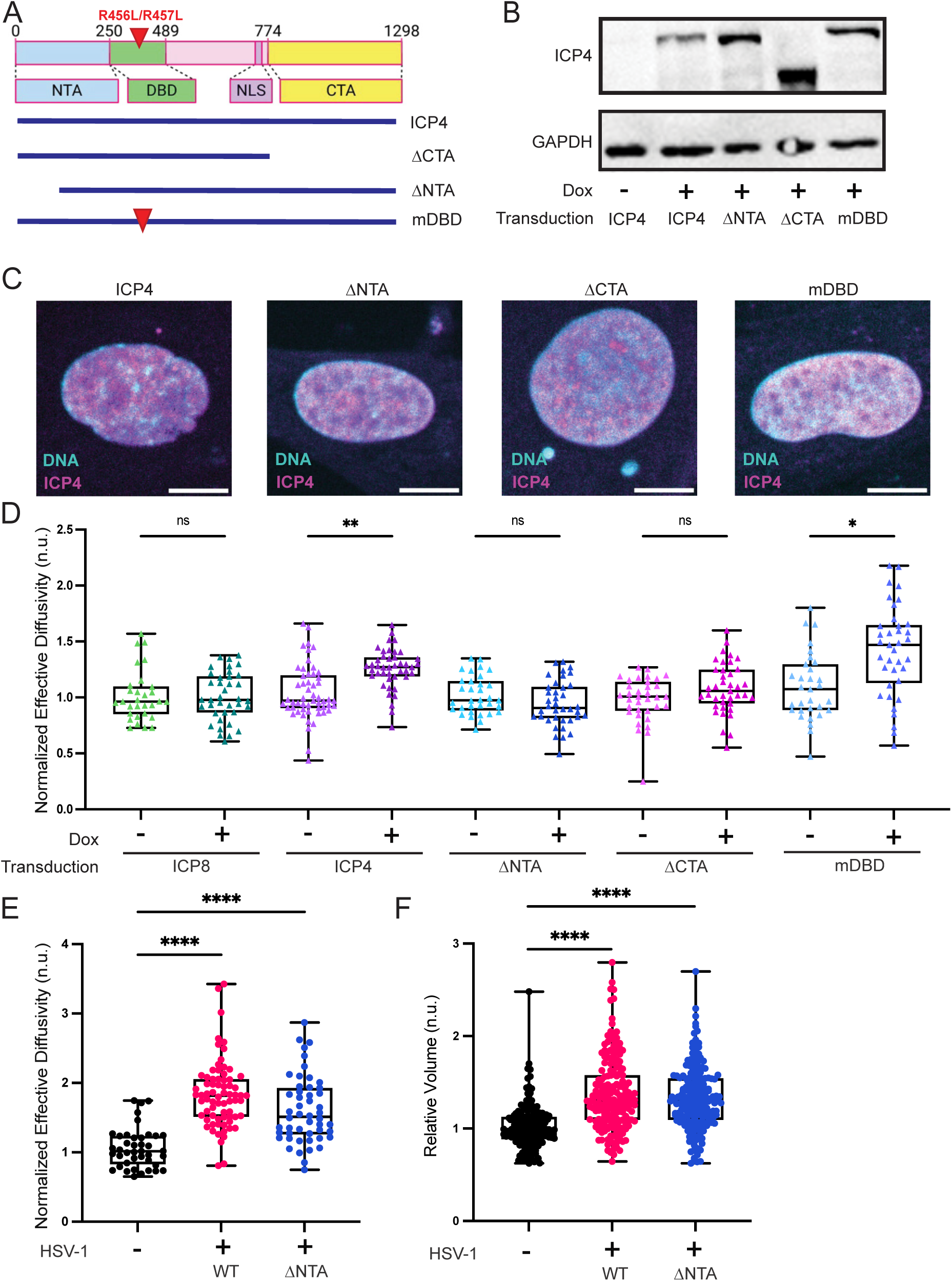
Sequences at both the N- and C-terminus of ICP4 are required to increase nucGEM diffusivity in uninfected cells. (A) Schematic of ICP4 domains and mutants. (B) Immunoblot of ICP4 expression for the HSV-1 mutants used in (A). (C) IF of exogenous ICP4. Scale bar, 10 μm. (D) Effective diffusivity (D_eff_) of nucGEMs in NHDFs transduced with lentiviruses expressing codon-optimized ICP8, ICP4, or ICP4 mutants illustrated in (A) under a tet-promoter and induced with doxycycline for 9 h. n>113; N≥5 biological replicates. (E) Effective diffusivity of nucGEMs in NHDFs infected with WT HSV-1 or ΔNTA mutant HSV-1 at 9 hpi, MOI = 5. n>116; N≥3 biological replicates. (F) Relative nuclear volume of cells from (E), calculated with Foci-Counting. All volumes were normalized to the median uninfected nuclear volume by experiment. n>357; N≥3 biological replicates. IF images and immunoblots are representative of N≥3 biological replicates. *p<0.05, **p<0.01, ****p<0.0001 by Kruskal-Wallis test.

A ΔNTA mutant virus (producing 10-fold lower titer than WT)^64^ partially increased nucGEM diffusivity **(Fig. 4E)**, indicating that this domain is not fully required for fluidization in the presence of other virus activities. The ΔNTA mutant virus drove an increase in nuclear volume (**Fig. 4F**), suggesting an additional ICP4-independent mechanism for nuclear fluidization related to nuclear volume increase in the context of full virus infection. Thus, ICP4-driven fluidization requires NTA and CTA but not specific DNA binding.

### ICP4 induction does not alter global transcription

Since transcription increases non-thermal energy and crowding, and ICP4 is a transcription factor^67^, we tested whether ICP4 induction alters global transcription. We incubated cells for 2 h with the nucleoside analog 5-ethynyl uridine (5EU). 5EU is incorporated into newly synthesized RNA and can subsequently be visualized by fixing cells and using click chemistry to conjugate its alkyne group to an azide-modified fluorescent dye^68,69^. The intensity of fluorescence is proportional to the total amount of RNA produced during the pulse of 5EU incubation. Neither ICP4, ICP8, ICP4-ΔNTA, nor ICP4-ΔCTA induction changed 5EU incorporation (**Fig. S4B**). Thus, nuclear fluidization by exogenous ICP4 is unlikely to be due to global changes in host cell transcription.

### ICP4 induction increases motion of bound histones

The biophysical properties of the nucleus are also affected by chromatin—confinement, porosity, and non-thermal motion can all be affected by chromatin organization and active processes including chromatin remodeling^70–72^. Previous work on ICP4 showed that it co-purifies with the chromatin remodeling complexes SWI/SNF, INO80, and NuRD^73^. To determine if increased chromatin dynamics could explain the ICP4-driven increase in nucGEM diffusivity, we transduced primary NHDFs with a HaloTag-fused histone 2A (Halo-H2A) construct. HaloTag is a self-labeling tag that can bind covalently to bright fluorescent HaloTag Ligands (HTLs) enabling SPT^74^. We tracked Halo-H2A by highly inclined and laminated optical sheet (HILO) microscopy at 40 Hz (capturing bound and unbound histones) and 4 Hz (bound only, as free histones blur at low frame rates; **Fig. 5A**). At 40 Hz, only WT infection increased Halo-H2A diffusivity (**Fig. 5B**). At 4 Hz (bound histones), both WT and ΔICP8 increased diffusivity (**Fig. 5C**). TPM revealed correlated histone motion up to 9 µm in uninfected cells (Fig. 5D, Fig. S**5A-B**). HSV-1 infection decreased most correlations but slightly increased correlation at 1.8 µm (**Fig. 5D**). Chromatin elasticity and viscosity decreased during infection (**Fig. S5C**), matching nucGEM results and confirming HSV-1 increases chromatin motion independently of DNA replication.

**Figure 5.**
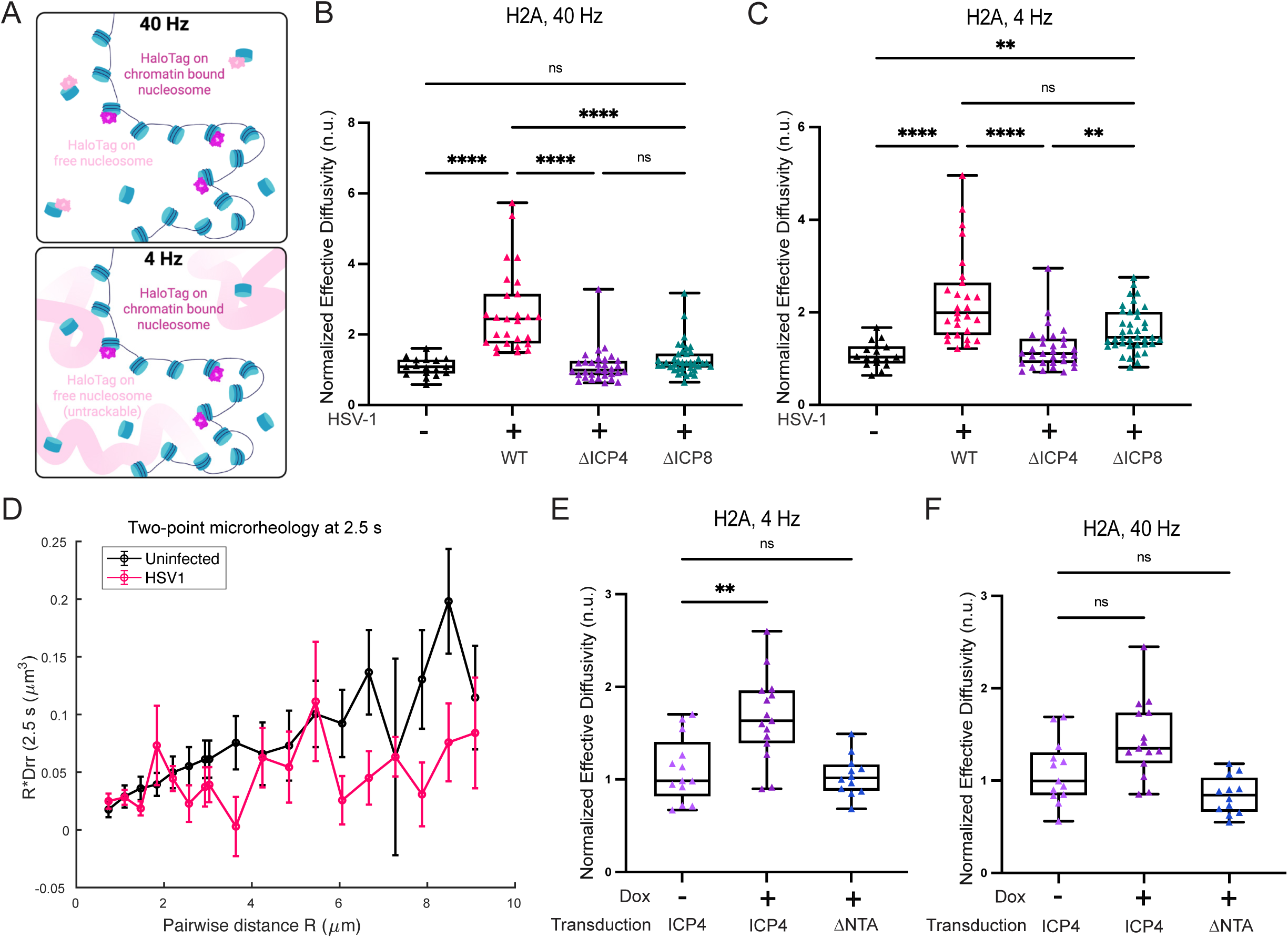
Movement of bound histones increases during ICP4 induction and during HSV-1 infection dependent on the presence of ICP4. (A) Schematic showing single-particle tracking (SPT) of tagged histones at short (40 Hz) and long (4 Hz) time scales. Light pink particles represent freely diffusing histones, which are only visible at 40 Hz. Dark pink particles represent bound histones, which are visible at 40 Hz and 4 Hz. (B, C) NHDFs expressing Halo-H2A were infected with WT HSV- 1, ΔICP4, or ΔICP8 and SPT was performed at 9 hpi to capture freely diffusing (B, 40 Hz) and bound (C, 4 Hz) histones. n>45; N≥3 biological replicates. (D) TPM at 2.5 s quantifying the correlated motion for histone pairs separated by a distance range [R] in NHDFs +/- HSV-1 identified in (C). (E, F) Effective diffusivity at 4 Hz (E) and 40 Hz (F) of Halo-H2A in NHDFs transduced with lentiviruses expressing either ICP4 or ICP4-ΔNTA under a tet-promoter and induced with doxycycline for 9 h. n>35; N≥3 biological replicates. **p<0.01, ****p<0.0001 by Kruskal-Wallis test.

We then assessed whether ICP4 induction alone was sufficient to increase Halo-H2A motion. ICP4 induction increased bound H2A motion (**Fig. 5E**) and decreased chromatin elasticity and viscosity (**Fig. S5D**). ICP4-ΔNTA did not increase H2A motion at either timescale (**Fig. 5E-F**), consistent with nucGEM data.

### High cell confluency and osmotic compression attenuate nuclear fluidization

To ascertain the importance of nuclear fluidization for HSV-1 infection, we used two orthogonal approaches to alter the biophysical properties of the nucleus. First, we grew cells to high confluence. We previously found that, upon contact inhibition, epithelial cells stop growing but undergo an additional cell division resulting in decreased cell and nuclear volume^75^. We confirmed that high confluence also decreased nuclear volume in NHDFs (**Fig. S6A**). Decreased nuclear volume is predicted to increase the concentration of macromolecules and chromatin. Consistent with this prediction, nucGEM diffusivity was significantly decreased in these confluent cells (**Fig. S6B**). Furthermore, the increase in nucGEM diffusivity during HSV-1 infection was attenuated in these conditions (**Fig. 6A, Fig. S6C-D**). As an orthogonal approach, we used osmotic compression. 150mM sorbitol decreased nucGEM diffusivity sufficiently to mostly prevent fluidization at 9 hpi (Fig 6A, Fig. S**6B-C,E**).

**Figure 6.**
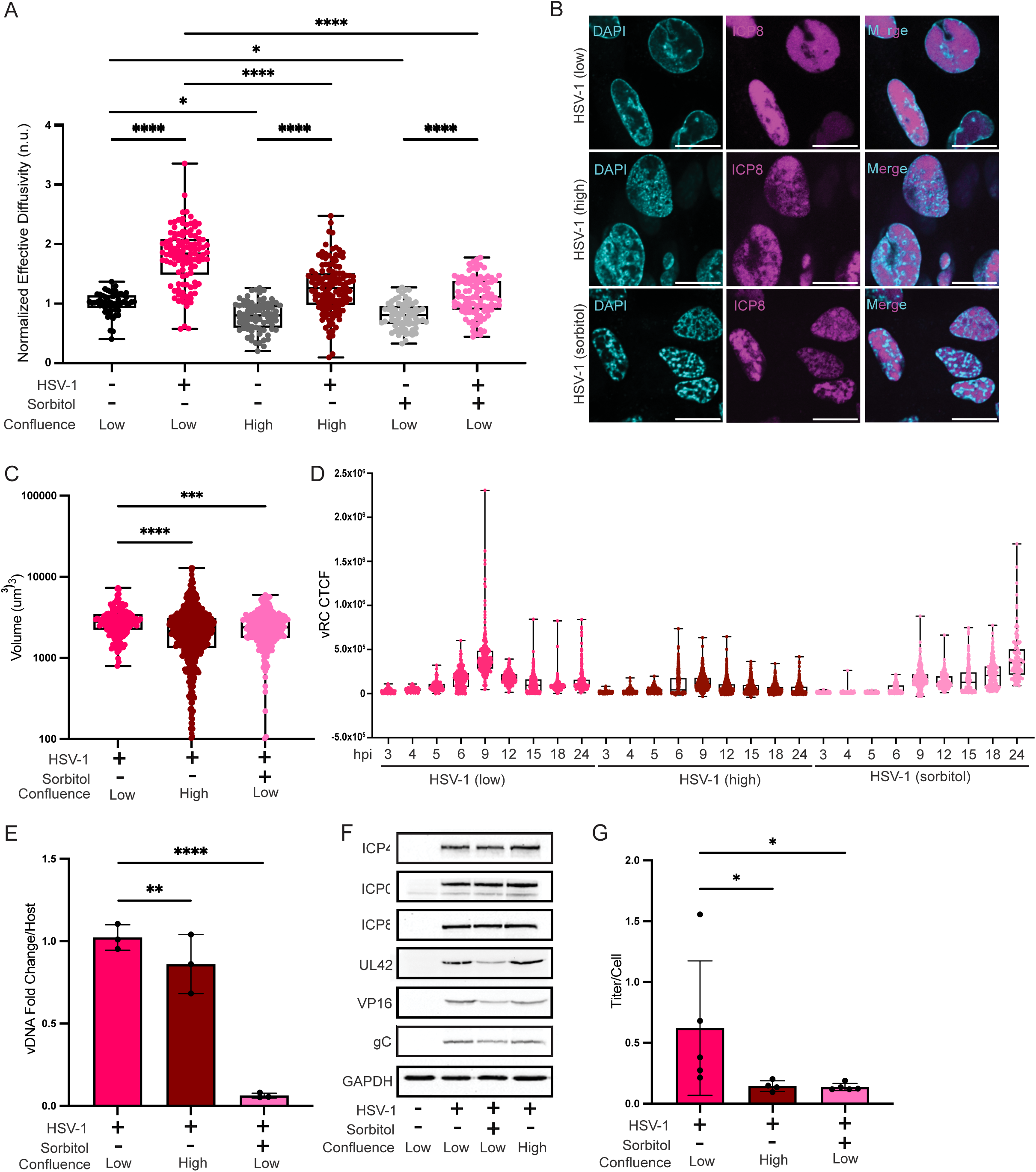
Attenuating nuclear fluidization disrupts viral replication compartment formation and decreases infectious virus production. (A) Effective diffusivity of nucGEMs in NHDFs grown to confluence (high) or under osmotic pressure (sorbitol) compared to NHDFs at 70% confluence (low), +/- WT HSV-1 at 9hpi, MOI = 5. n>136; N≥3 biological replicates. (B) Indirect IF for cells from (A) to detect ICP8, a marker of viral replication compartments (vRCs). Scale bar, 20 μm. (C) Total vRC volume from cells in (A). n>784; N≥4 biological replicates. (D) NHDFs were treated as in (A) and fixed for IF as in (B) at the indicated time points. Corrected total cell fluorescence (CTCF) from ICP8 indirect IF detection. n>305; N≥3 biological replicates. Statistical analysis available in **Table 2**. (E) Total DNA was collected at 9 hpi and analyzed by qPCR to quantify total copies of viral DNA (fold change over HSV-1 (low).) N≥3 biological replicates. (F) Immunoblot for a representative panel of IE, E, and L HSV-1 proteins in HSV-1 (high) and HSV-1 (sorbitol) compared to HSV-1 (low). (G) Supernatant was collected at 9 hpi and quantified by plaque assay to determine infectious titer per cell. N≥4 biological replicates. IF images are representative of N≥3 biological replicates. *p<0.05, **p<0.01, ***p<0.001, ****p<0.0001 by two-way ANOVA with Bonferroni’s correction (A), by one-way ANOVA (C,G), or by linear mixed-effects model (E).

### Preventing nuclear fluidization disrupts viral replication compartment maturation and decreases infectious virus production

The formation of viral replication compartments (vRCs) is a key step in progression of the HSV-1 life cycle^76,77^. Formation of vRCs has been shown to proceed via formation of pre-vRCs and their subsequent coalescence into vRCs^31,32^. Since fluidity affects condensate formation^12,13^, we hypothesized nuclear fluidization enables vRC coalescence.

To test this idea, we infected cells with HSV-1 in control, high confluence, and sorbitol conditions, and probed for ICP8 (a vRC marker). Indeed, when nuclear fluidization was prevented by high confluence or sorbitol treatment, vRCs did not fully mature into a single vRC, but instead multiple smaller vRCs formed (**Fig. 6B**), reducing total vRC volume (**Fig. 6C**).

To determine if the differences in vRC maturation were due to delayed life cycle progression under high confluence and sorbitol conditions, we performed a time course assaying vRC maturation under all three conditions from 3 hpi to 24 hpi. While the maturation of vRCs appeared to be delayed under sorbitol treatment compared to untreated cells, vRCs did not detectably coalesce in sorbitol treated or high confluence conditions (**Fig. 6D, Fig. S6F-G**) even at much later timepoints. These data suggest that nuclear fluidization enables vRC coalescence

To assess the function of vRCs, we measured viral DNA replication using qPCR under high confluence and sorbitol treatment compared to WT HSV-1 infection at 9 hpi. Sorbitol treatment led to a ten-fold reduction in viral DNA copy number, while high confluence led to a modest, but significant reduction of viral DNA copy number **(Fig. 6E).**

We also assessed the production of virus proteins. High confluence did not impact the accumulation of representative virus IE, E or L proteins compared to infection of sub-confluent cells, and sorbitol treatment only partially impacted virus protein accumulation (**Fig. 6F**).

Finally, we collected the supernatant from each condition and performed plaque assays to calculate the infectious titer. Both high confluence and sorbitol treatment reduced viral titer by approximately three-fold and four-fold, respectively (**Fig. 6G)**.

Together, these results are consistent with the hypothesis that nuclear fluidization enables vRC condensate formation and optimal progression of the HSV-1 life cycle.

### Fluidization of the nucleus by ICP4 facilitates growth of synthetic condensates

Natural condensates are complex, and so it can be difficult to interpret changes in their structure. Therefore, we used a synthetic system to determine how nuclear fluidization by ICP4 impacts condensate formation. Here, a chemical dimerization system initiates multivalent interactions between a fluorescently-tagged coiled-coil protein and nuclear oligomer (**Fig. 7A**, methods)^15^. This system was previously shown to grow mostly through a nucleation and coalescence mechanism^15^, similar to vRCs^32^.

**Figure 7.**
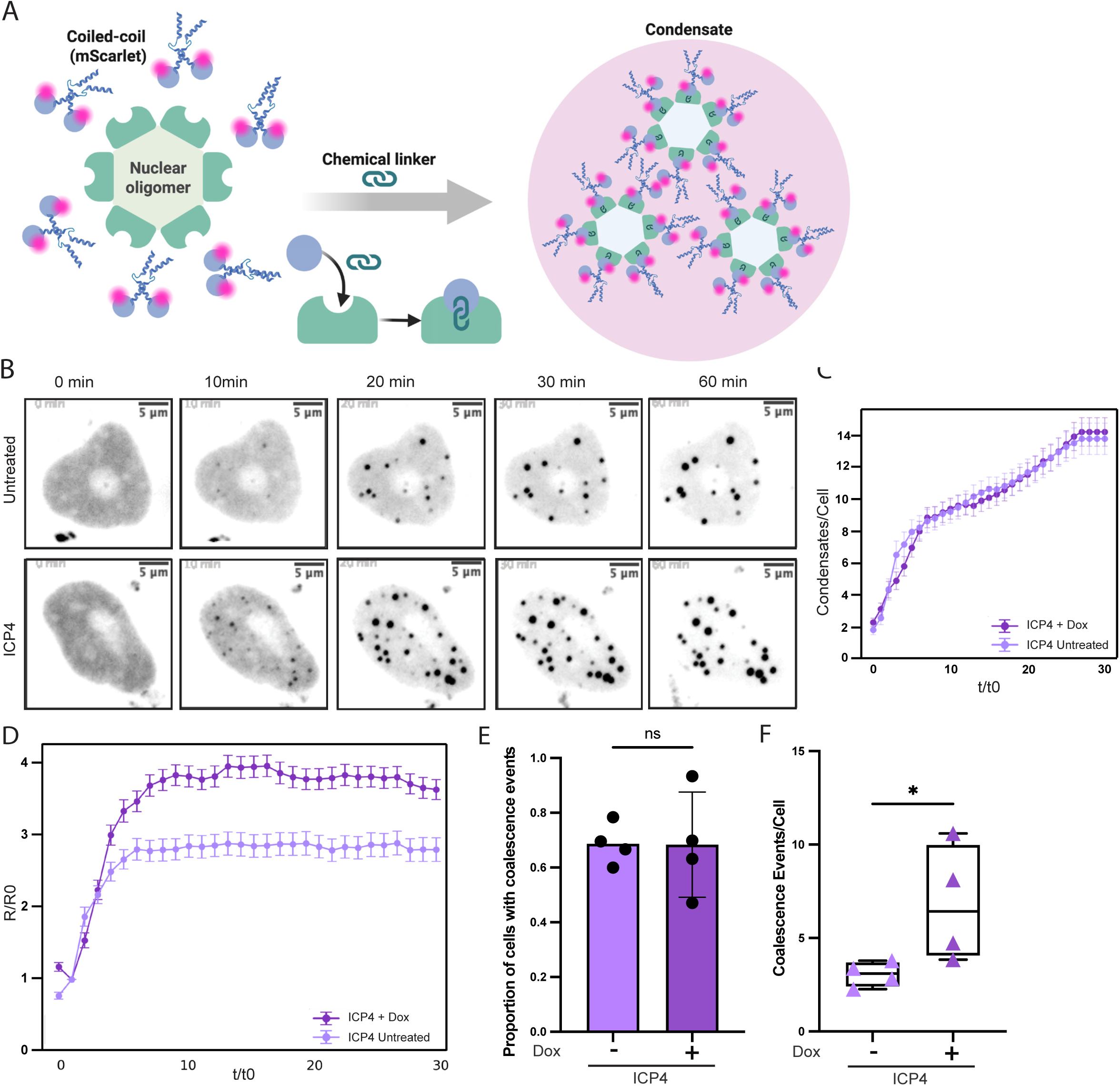
Fluidization by ICP4 facilitates artificial condensate growth. (A) Schematic of a two-component artificial condensate system. One component is a fluorescently-tagged coiled-coil phase-separating protein (Mad1, blue helices) fused to E. coli dihydrofolate reductase (eDHFR, blue circles.) The second component is a nuclear hexamer (HO Tag3, light green) fused to HaloTag (dark green). The TFH ligand causes dimerization of eDFHR and Halo, forming condensates that are visualizable through fluorescence microscopy. (B) ICP4 was induced for 8 h, or cells were left untreated. Cells were imaged every 2 minutes after induction of condensate formation by TFH treatment for a total of 1 h. Images show max projections of nuclear condensates. Scale bar, 5 μm. (C) Average number of condensates identified per cell with (+ Dox) or without (untreated) ICP4 expression. Time was normalized to the first timepoint at which condensates nucleated (t/t0). n>23; N≥3 biological replicates. (D) Average condensate radii over time in cells from (C). Radii were normalized to the initial droplet size at nucleation (R/R0). (E) Proportion of cells with (+) or without (-) ICP4 expression exhibiting condensate coalescence events. n>135; N≥4 biological replicates. (F) Average number of coalescence events per cell in cells from (E) with (+) or without (-) ICP4 expression. n>135; N≥4 biological replicates. Statistical analysis for (C,D) available in **Table 3**. *p<0.05 by Mann-Whitney test.

We transfected HeLa cells with plasmids encoding inducible ICP4 and the two components of the synthetic condensates. We initiated condensate formation by addition of the chemical dimerizer, either after 8h ICP4 induction to drive nuclear fluidization or in control cells, and observed condensate formation kinetics for 1 h (**Fig. 7B**). We blotted for the components in each condition and determined there was no difference in condensate component production during ICP4 induction compared to uninduced cells **(Supplementary Fig. 7A).** There was no difference in the average number of condensates per cell across conditions **(Fig. 7C),** indicating that there was no significant difference in nucleation between the two conditions. However, condensate growth was more efficient after ICP4 induction, as measured by average condensate radius **(Fig. 7D**) and average condensate intensity (**Fig. S7B**). These results indicate that fluidization of the nucleus by ICP4 induction is sufficient to facilitate condensate growth.

We next analyzed condensate kinetics for 3 h following TFH treatment and found that, while the proportion of cells exhibiting coalescence events was comparable between conditions (**Fig. 7E**), ICP4 expression led to an increase in the average number of coalescence events per cell (**Fig. 7F**). ICP4 induction increased the effective diffusivity and anomalous exponent of these condensates, which we estimate to be 300 - 1200 nm, consistent with fluidization at this larger scale, (**Fig. S7C-D**). Taken together, these results show that ICP4 induction fluidizes the nucleus, enabling condensate growth by coalescence.

## DISCUSSION

The mechanisms that define the physical parameters of the nucleus are poorly understood, perhaps because major perturbations would be highly deleterious. We reasoned that nuclear viruses might encode activities that acutely perturb these properties and thereby provide a path to understand the control of this crucial parameter in normal biology.

We discovered a dramatic increase in mesoscale diffusivity in the nucleus during HSV-1 infection that is linked to an increase in chromatin dynamics, independent of previously described architectural changes. We confirmed that this increase in diffusivity was due to fluidization of the nucleus by using probe particles of varying sizes (12, 40 and 50 nm nanoparticles, as well as condensates in the 300 - 1200 nm diameter range) as well as analyses of the storage modulus (elasticity) and loss modulus (viscosity) to correlate increases in GEM motion with a changing viscoelastic environment. We observed reductions in both elasticity and viscosity during HSV-1 infection and ICP4 induction compared to control conditions. Furthermore, two point microrheology suggested that HSV-1 infection increased non-thermal activity in the nucleus. A single factor, the immediate early transcription factor ICP4, drives this nuclear fluidization. This fluidization was important for condensate growth, which has consequences for the maturation of viral replication compartments (vRCs) and thus HSV-1 life cycle progression.

HSV-1 infection was known to have global effects on nuclear size and chromatin structure: host chromatin is marginated leading to enlargement of the interchromatin space^20^, peripheral chromatin restructuring occurs late in infection to allow for virus egress^40,59^, and nuclear volume is increased^20^. However, these changes are all dependent on virus DNA replication^41^, while the increased mesoscale fluidity that we discovered is independent of these processes, indicating a distinct mechanism. Furthermore, nuclear fluidization from a DNA-replication deficient HSV-1 virus or exogenous ICP4 expression both occurred without margination of host chromatin or increased nuclear volume.

Although we have yet to elucidate the detailed mechanism by which HSV-1 increases nuclear fluidity, there are several promising possibilities. Chromatin creates significant elastic confinement, reducing mesoscale particle motion^78^ and frustrating mesoscale condensate growth^14^. Reducing this elastic confinement is one way to increase fluidity. Reduced chromatin confinement can be achieved through an increase in non-thermal motion, for example from increased transcription or increased chromatin remodeling^12,79–81^. ICP4 binds promiscuously to host chromatin early in infection, and recruits both the transcriptional machinery and chromatin remodelers including the ATPase subunits of the SWI/SNF, INO80, and NuRD complexes^73,82^. We did not detect changes in global transcription rates during ICP4 induction, but we did observe increased motion of bound Halo-H2A, consistent with increased chromatin motion. Additionally, the decrease in storage modulus seen during HSV-1 infection and ICP4 induction in both nucGEMs and bound Halo-H2A supports the hypothesis that HSV-1 infection, and ICP4 expression in particular, lead to a reduction in elastic confinement. Recent studies suggest that SWI/SNF activity decreases the global stiffness of the nucleus^80^, and NuRD activity increases chromatin motion^81^. Further work will be required to understand the precise effects of ICP4 on SWI/SNF, INO80, and NuRD; it is interesting that a single viral factor may modulate multiple host chromatin remodeling complexes. Overall, prior literature and our observations are consistent with a model where ICP4 fluidizes the nucleus by altering chromatin dynamics.

The hypothesis that fluidization is mediated, at least in part, by changes in chromatin dynamics is further supported by two-point microrheology (TPM) data extracted from nucGEM and Halo-H2A trajectory analysis. We found correlated motion of bound Halo-H2A in uninfected cells at length scales up to 9 µm. These observations are consistent with previous reports from displacement correlation spectroscopy (DCS) measurements^83,84^. These long-range correlations were attenuated during HSV-1 infection. DCS studies found that perturbation of nuclear ATPases such as DNA polymerase, RNA polymerase II, and topoisomerase II can disrupt large-scale coherent chromatin motion while simultaneously increasing local movement^83^. We therefore hypothesize that ICP4 fluidizes the nucleus by modulating host chromatin remodeling complexes, leading to disruption of larger chromatin domains but increasing short-range chromatin motion.

Our TPM on nucGEMs suggests a model whereby correlated motion between chromatin compartments contributes to particle diffusivity in the nucleoplasm. Specifically, there is no correlated motion at short (0.7 µm) or long (>1.8 µm) distances, but clear correlated motion at 1.1 to 1.47 µm separation in uninfected cells. This separation is consistent with the distance between interchromatin compartments^85,86^. Thus, we speculate that nucGEMs < 1 µm apart move independently of one another within the same chromatin compartment, while nucGEMs at 1.1 to 1.47 µm separation exhibit correlated motion because adjacent interchromatin compartments move together.

HSV-1 infection increased the correlated motion of nucGEMs at the 1.5 to 2.5 µm length-scale, consistent with a model where HSV-1 infection drives nuclear fluidization. It is additionally known that chromatin motion is locally correlated in uninfected cells^83,84,87^, a finding that we have verified using TPM across a range of distances. However, this correlation of chromatin motion is largely disrupted during HSV-1 infection, particularly at longer length scales, perhaps due to chromatin margination, which likely disrupts chromatin domain organization^20,41,88^. We hypothesize that the resulting increase in nucleoplasmic homogeneity, together with reduced viscosity and elasticity allows active fluctuations to propagate more freely in this enlarged interchromosomal space, thereby enhancing the correlated motion observed between nucGEM pairs.

Taking all of these observations together, we speculate that ICP4 modulates some combination of the SWI/SNF, INO80, and NuRD complexes leading to an increase in mesoscale effective energy, a decrease in the effective persistence length of chromatin, and a decrease in elastic confinement. Future work will test these ideas through further structural and dynamical analyses.

HSV-1 might alter the biophysical properties of the nucleus to inhibit host functions or enable viral processes. We speculate that the low mesoscale fluidity in nuclei could be an intrinsic barrier to the replication of all nuclear viruses. This could be particularly relevant in the nuclei of fibroblasts and other epithelial cells resident within dense tissues, where lytic HSV-1 replication typically occurs. Consistent with this idea, vRC formation and virus production was inefficient in high-density cell cultures that had low mesoscale fluidity. Nuclear fluidization by ICP4 might help HSV-1 overcome this fundamental physical barrier to replication.

Condensate formation is highly sensitive to the physical properties of the cell interior^12,13^. In particular, chromatin has been shown to mechanically frustrate growth of large condensates in the nucleus, while fluidization of the cellular environment through increased fluidity, reduced confinement, or increased non-thermal energy has been shown to promote droplet growth^13,14^. Using a synthetic system, we confirmed that ICP4 expression is sufficient to increase the efficiency of nuclear condensate formation. We also showed that condensate coalescence increased during ICP4-driven fluidization, while preventing nuclear fluidization frustrated HSV-1 vRC growth and maturation. These data suggest that HSV-1 fluidizes the nucleus to enable the growth, maturation, and efficient function of vRCs, thus driving progression of the virus life cycle. It will be interesting to examine whether this increase in nuclear fluidization is specific to HSV-1, or is conserved across the herpesvirus family, dsDNA viruses, or beyond.

Little is known about the control of mesoscale nuclear physical properties and their effects on biology. This is partially due to the difficulty of perturbing these properties without losing viability. This issue can be surmounted by studying lytic viruses, an approach that has led to the discovery of fundamental mechanisms of cellular control. Studying sarcoma viruses led to the discovery of the Ras^89,90^ and Src oncogenes^91–93^ that control cell division initiation, while studying adenovirus led to the discovery of p53, which controls apoptosis^94–97^. We have determined that a single HSV-1 protein, ICP4, increases nuclear fluidity. Further mechanistic study of ICP4 provides a path to discover mechanisms by which the physical properties of the nucleus are controlled, and to understand how these properties influence both cellular and viral function.

### Limitations of the Study

We have not identified the specific molecular mechanism by which ICP4 drives nuclear fluidization, though previous studies suggest that interactions with chromatin remodeling complexes are a promising direction for future investigation. Understanding the contribution of active processes to the fluidization phenomenon is of great interest, but beyond the scope of this study. We were unable to visualize vRC formation by live imaging, but recent development of additional tools^98^ provides an opportunity for future studies. The difficulty in obtaining three dimensional data at high frame-rates limits our ability to analyze nucGEM diffusivity at long timescales, which prevents access of D(∞) and thus analysis of elastic recoil that could further parse our tracking data.

## RESOURCE AVAILABILITY

### Lead Contact

Requests for further information and resources should be directed to and will be fulfilled by the lead contact, Liam J. Holt.

### Materials Availability

Plasmids generated in this study have been deposited to Addgene. All materials will be made available on request.

### Data and Code Availability

● All original code has been deposited on GitHub (https://github.com/nlherzog/Biophysical-Tracking)
● Original imaging data have been deposited at Mendeley Data (doi: 10.17632/tmbs5mpcp3.1) and are publicly available as of the date of publication.
● Any additional information required to reanalyze the data reported in this paper is available from the lead contact upon request.

## ACKNOWLEDGEMENTS

LJH was supported by the Hypothesis Fund https://doi.org/10.13039/100031064 and National Institutes of Health (NIH) grants GM132447 and AG090618; AW by NIH grants AI176335 and AI170583; IM by NIH grants GM056927, AI073898, and AI176335; HZ by the National Science Foundation (NSF MCB-2145083).

We thank Eli Rothenberg and Tafadzwa Chigumira for advice on experimental design and data interpretation, Andrew Bazley for assistance with microscopy, and David Duran-Chaparro for analysis codes. Neal DeLuca, Priscilla Schaffer, Steve Rice, David Knipe, Eli Rothenberg, and Jens Bosse generously supplied viruses, cell lines, and other experimental tools. We thank the Lavis lab and Open Chemistry team (HHMI Janelia Research Campus) for the gift of JF and JFX dyes. Lastly, we thank all members of the Holt, Mohr, and Wilson labs for their feedback, advice, and encouragement.

## AUTHOR CONTRIBUTIONS

N.L.H. designed and conducted experiments with help from G.R.K and F.K. N.L.H and T.S. performed data analysis with help from S.K. T.S. performed all two-point microrheology analysis and analyzed the complex modulus. D.M.C created the reagents for the inducible artificial condensate system, supervised by H.Z. All aspects of the study were supervised by I.M., A.C.W., and L.J.H. N.L.H., T.S., G.R.K., I.M., A.C.W., and L.J.H. contributed to writing and editing the manuscript.

## DECLARATION OF INTERESTS

The authors declare no competing interests.

## STAR METHODS

### EXPERIMENTAL MODEL AND STUDY PARTICIPANT DETAILS

#### Human Cell Lines

Normal human dermal fibroblasts (Lonza; CC-2509), HeLa (Jef Boeke Lab donation), and HEK293T cells (Jef Boeke Lab donation) were cultured in Dulbecco’s modification of Eagle’s medium (DMEM) (Corning; 10-013-CV) supplemented with 10% (v/v) fetal bovine serum (FBS) (Gemini bio-products; 100-106) and 100 U/mL penicillin-100 μg/mL streptomycin (Corning; MT-30-002-Cl). U2OS cells (ATCC; HTB-96), Vero cells (ATCC; CCL-81), V22 cells, V529 cells, and E5 cells were cultured in DMEM supplemented with 5% (v/v) FBS and 100 U/mL penicillin-100 μg/mL streptomycin. V27 cells were cultured in DMEM supplemented with 5% (v/v) fetal bovine serum (FBS) and 100 U/mL penicillin-100 μg/mL streptomycin under selection with 400 µg/mL of G418 (Sigma-Aldrich; 04727878001). All cells were grown in a humidified incubator atmosphere at 37°C and 5% CO2.

### METHOD DETAILS

#### Virus growth and infection

Wild-type HSV-1 KOS was grown on Vero cells. ΔICP4 HSV-1 was grown on E5 cells, both donated by Neal DeLuca (University of Pittsburgh). ΔICP0 HSV-1 was grown on U2OS cells. ΔICP8 HSV-1 was grown on V529 cells, both provided by David Knipe (Harvard Medical School). ΔICP22 HSV-1 was grown on V22 cells; ΔICP27 HSV-1 was grown on V27 cells, all obtained from Steve Rice (University of Minnesota). UV-inactivated virus was prepared by exposing 1 mL layers of virus stock in a six-well dish to six pulses of 0.12 J/cm^2^ of UV light in a Stratalinker (Stratagene). To grow virus stocks, cells were grown to 70-80% confluency in 10 cm dishes, then infected at an MOI of 0.001 in DMEM supplemented with 1% FBS. After 48-72 hours, cells and supernatants were collected and subjected to three freeze-thaw cycles before being aliquoted and stored in -80°C for future use. Virus stock titers were determined using a plaque assay on Vero cells or the appropriate complementing Vero-derived cell line. For all infection experiments, cells were seeded on a 24-well (Cellvis; P24-1.5H-N) or 12-well (Cellvis; P12-1.5H-N) glass bottom plate 24 hours prior to experiment start. Cells were infected at MOI=5 in DMEM with 1% FBS for all virus strains except ΔICP0, which was used at MOI = 10. Media was replaced after 1.5 h of incubation to full media (DMEM + 10% FBS + 1% P/S). Cells were imaged, total protein was harvested for immunoblotting, total RNA was harvested for RNA extraction, or cells were fixed for immunofluorescence at indicated timepoints. For experiments under confluent conditions, cells were seeded at 100% confluency 48 h prior to the start of the experiment. Cells were infected with MOI = 5. MOI was calculated based on hemocytometer counts of wells at 70-80% confluence and 100% confluence, respectively.

#### Lentivirus/Retrovirus creation

HEK293T cells (9 × 10^6^ per 15 cm dish) were plated in antibiotic-free DMEM (Gibco, Cat. No. 11995073) supplemented with 10% FBS. The next day, cells were transfected with appropriate transgene plasmid together with packaging plasmids (psPAX2 and pMD2.G for lentivirus, VSV-G and GP for retrovirus), using fuGENE HD transfection reagent (Promega, Cat. no. E2312) following manufacturer’s protocol. 24 h later, 15 mL antibiotic-free DMEM was replaced, and the supernatants were collected at 72 h post-transfection, aliquoted and stored at -80°C until later use. Both lentivirus and retrovirus were introduced into NHDF wild type and NHDF-nucPfV cell lines of interest via reverse transduction with 5 mL of virus in fresh media in the presence of 10 µg/mL polybrene (Sigma-Aldrich; TR-1003-G), replacing media after 24 h. Cells were selected with puromycin for three days. 24 h prior to experiment start, cells were split to a 24-well (Cellvis; P24-1.5H-N) or 12-well (Cellvis; P12-1.5H-N) glass bottom plate. Cells were freshly transduced with lentivirus for each new experiment and used within 2 passages of transduction. For cells transduced with retrovirus, after selecting antibiotic resistant cell populations, they were frozen in 10% DMSO (Sigma-Aldrich, Cat. no. D2650- 100) in FBS and thawed for use in experiments whenever needed.

#### Plasmid construction

Sequence blocks encoding codon optimized HSV-1 ICP4 and ICP8 (based on KOS strain) were obtained from Integrated DNA technologies. The custom vector for a Tet-inducible ICP4 expression plasmid was from VectorBuilder (VB211009-1101fac: https://en.vectorbuilder.com/). This plasmid was used directly for transfection in HeLa cells. A codon-optimized ICP8 custom gene block was purchased from Twist Bioscience (https://ecommerce.twistdna.com/). To construct lentiviral vectors expressing ICP4 and ICP8, respectively, the assembled coding sequences were incorporated into a Tet-inducible lentivirus vector via Gibson assembly.

To generate ICP4 mutant proteins, primers were generated to amplify segments lacking either the C-terminal activation domain or the first 90 amino acids of the N-terminal activation domain of ICP4 from the codon optimized ICP4 present in pLH2488^99,100^. pLH2631 and pLH2632 were made using Gibson assembly of the PCR DNA fragments encoding the KOZAK sequence and ICP4-ΔNTA or ICP4-ΔCTA respectively. The pLVX-TetOne-puro plasmid backbone was digested from pLH2537 using EcoRI and BamHI, then purified and ligated to the ICP4-ΔNTA and ICP4-ΔCTA fragments. Localized mutations in the DNA-binding domain of ICP4 (ICP4-*m*DBD-RR/LL and ICP4-*m*DBD-KG/NC) were introduced into pLH2488 using overlapping primers containing the mutation of interest to generate two PCR products that had homology to one another^65^. The products were then assembled using NEB HiFi DNA assembly to generate pLH2658 and pLH2659. All constructs were confirmed by DNA sequencing.

To generate the plasmid encoding the 12nm-nucGEMs, we first synthesized the gene block containing the de novo-designed monomer (PDB ID: 8F53)^49^ followed by a 4X(GGGGS) linker and a mammalian codon-optimized T-Sapphire fluorescent protein (Twist Bioscience). To increase capsid assembly efficiency, the HOTag3 (homo-oligomeric tag3)^101^ was added at the N-terminus of the monomer. An SV40 nuclear localization signal (SV40NLS) was then appended to the N-terminus of HOTag3 to direct the assembled particles to the nucleus. The final construct therefore consisted of SV40NLS-HOTag3-monomer-4X(GGGGS)-TSapphire. Each gene fragment was PCR-amplified using primers containing 25bp overlaps with adjacent fragments. The resulting components were then assembled into a lentiviral plasmid under the UBC promoter using Gibson assembly to generate pLH2899, and the construct was verified by DNA sequencing.

#### Transient transfection

For nucGEM experiments in HeLa cells, cells were seeded at 60%–70% confluency in a 6-well tissue culture plate (CELLTREAT; 229106) and transfected the next day with 1 μg of plasmid DNA (pLH2112) per well using fuGENE HD reagent per manufacturer guidelines. For artificial condensate experiments, 6 x 10^5^ HeLa cells were reverse transfected in a 6-well tissue culture plate (CELLTREAT; 229106) with 0.8 µg of ICP4 expressing plasmid (pLH2112) and 0.5 µg of each plasmid encoding an artificial condensate component (pLH2712 and pLH2714). pLH2712 contains the nuclear hexamer HO Tag3 fused to HaloTag. pLH2714 contains the phase-separating coiled-coil protein Mad1 with an N-terminal fusion to mScarlet-eDHFR^102^ to facilitate visualization and binding to the HaloTag of pLH2712 upon chemical dimerization. Culture media was replaced after 12 h and cells were split into a 24-well (Cellvis; P24-1.5H-N) or 12-well (Cellvis; P12-1.5H-N) glass bottom plate for imaging 24-48 h post-transfection. Imaging experiments were usually carried out between 48 and 72 h post-transfection.

#### Induction and chemical treatments

For all induction experiments, cells were seeded on a 24-well (Cellvis; P24-1.5H-N) or 12-well (Cellvis; P12-1.5H-N) glass bottom plate 24 hours prior to experiment start as previously described. For experiments blocking new protein synthesis, cells were pre-treated for 30 minutes with 15 µg/mL cycloheximide (CHX) (Sigma; C104450). CHX was refreshed during virus incubation, and again when virus inoculum was removed. For experiments blocking virus DNA replication, cells were treated with 300 µg/mL phosphonoacetic acid (PAA) (Sigma-Aldrich; 284270) beginning with virus addition. PAA was refreshed when virus inoculum was removed. To induce hyperosmotic stress, 150 mM sorbitol was added at 2.5 hpi, which roughly correlates with the onset of virus E protein expression as determined by immunoblotting. For doxycycline (dox) induction of ectopically expressed proteins, cells were treated with 3 µg/mL dox for 9 h prior to imaging or further processing. To perform single particle tracking of HaloTag-H2A (Halo-H2A), Halo-H2A was labeled with 10 pM JFX646 (gift of Janelia HHMI/Lavis Lab) in cell culture media for 30 minutes, then washed three times with PBS (Corning; 21-040-CV) before replacing with fresh media and proceeding to infection or induction as indicated.

#### Microscopy

For nucGEM experiments, artificial condensate assay experiments, as well as immunofluorescence imaging experiments, cells were mounted on a Nikon Ti2 X1 spinning disc confocal microscope and images were captured using a Prime 95B scMOS camera (Photometrics) with a 60x/1.49 numerical aperture objective lens with DIC capability. Fluorescence signals were obtained using 405 nm, 488 nm, 561 nm, and 640 nm lasers with the following emission filters: 455/50, 525/36, 605/52, and 705/72 (Chroma Technology Corp). For live cell imaging, the microscope was equipped with an incubator to maintain 37°C and 5% CO_2_. Cells were stained with a nuclear stain (SiR-DNA (Cytoskeleton; CY-SC007) or Hoechst (Thermo Fisher Scientific; 62249)) if applicable, then imaged at indicated timepoints. The nucGEMs were imaged at a rate of one image every 10 ms (100 Hz).

For single particle tracking of Halo-H2A, cells were mounted on a Nikon Ti Eclipse microscope equipped with a motorized Ti H-TIRF module and an Andor Zyla sCMOS camera behind a 0.7x relay magnification lens. Movies were acquired in highly inclined laminated optical sheet (HILO) mode as described in Tokunaga et al. ^103^. Data were collected using a CFI Apo 60× 1.49 NA TIRF objective and a quad-band STORM ultra-high S/N dichroic/emission filter (Nikon, 97335). Excitation was provided by a 640 nm laser (LUNF-XL, Nikon), and total internal reflection was calibrated using Nikon Elements software (insert version number). For HILO acquisition, the incident angle was manually adjusted to optimize Halo-H2A signal-to-noise, typically 5-10° below the critical angle and a 512×512 region of interest (ROI) was selected around the most evenly illuminated field of view. For our 250 ms timescale, we collected images at 25 ms exposure for 2,000 frames (40 Hz). To analyze data at 4 Hz (2500 ms timescale), max projections were created of every 10 slices to form a new collection of 200 total frames.

For live imaging of artificial condensate formation, the cells were mounted on a Nikon spinning disk confocal scanning microscope as previously described. Cells without condensates, but with high expression as determined by gating on the look up table (LUT) of both the anchor and the phase separating protein were selected and image acquisition began. At the end of the first time loop, an additional volume of growth media containing the chemical dimerizer TMP-Fluorobenzamide-Halo (TFH) to a final concentration of 50 μM was added to the stage to induce condensate formation in the mounted cells^102^. Z-stack images were collected with 1 μm spacing for a total depth of up to 12 μm, at 2 minute intervals for 1 hour or at 4 minute intervals for 4 hours.

#### Single particle tracking & analysis

Particles were identified and trajectories were generated using using FIJI plugin MOSAIC for ImageJ^104^. All trajectories from nucGEMs and Halo-H2A single particle tracking were analyzed with the GEMspa (GEM single particle analysis) software package that we are developing in house (https://github.com/liamholtlab/GEMspa, ^105^). Mean-square displacement (MSD) was calculated for every 2D trajectory, and only trajectories with more than 10 consecutive frames were used to calculate the time-averaged MSD, which was then fit to a linear model using the first 10 time intervals: MSD(**τ**) = 4D_eff_**τ**, where **τ** is the lag time (equal to the imaging frame interval) and D_eff_ is the effective diffusivity with the unit of μm^2^ / s. To determine the ensemble-time-averaged mean-square displacement (MSD) for each condition, all time-averaged MSDs were further averaged to fit with MSD(**τ**)τ_−ens_=4D**τ**^α^ where α is the anomalous exponent, with α = 1 being Brownian motion, α < 1 suggests sub-diffusive motion and α > 1 as super-diffusive motion. To generate region of interest (ROI) for marking individual cells, cellpose python package (https://github.com/mouseland/cellpose, ^106^) was used to segment based on the nuclear stain or the average projection intensity of Halo-H2A particles. These ROI were then input into GEMspa to quantify single cell effective diffusion. Cells were filtered based on density of identified trajectories and confirmed using GEMSpa Rainbow Tracks. All GEM analysis was additionally filtered for cells containing at least 10 trajectories of at least 10 time intervals in length. We used the median value of D_eff_ for single cell data to represent each condition. The median value of D_eff_ for each cell was normalized to the median value of D_eff_ in the control condition for each individual replicate of each experiment and this quantification was displayed using normalized units (n.u.) in each box and whiskers plot.

Normalized velocity autocorrelation was analyzed using a custom-developed MATLAB (R2019a) program^12^ based on experimental trajectory data^55^. Initially, velocities within specific time intervals were calculated for each trajectory, resulting in velocity time series along either the x or y directions. Subsequently, autocorrelation functions were applied to these velocity time series, generating outputs as a function of various time delays. Due to the orthogonality, the autocorrelation functions from both x and y direction were summed and were then normalized by the values at the zero time delay. For each condition, the normalized velocity autocorrelation functions were averaged across all trajectories.

To further characterize the viscoelastic properties of the nucleoplasm (from nucGEM tracks) and chromatin (from Halo-H2A tracks), we calculated the complex modulus *G*^∗^(*ω*) using the generalized Stokes-Einstein relation for viscoelastic materials^56,57^ given by the equations: 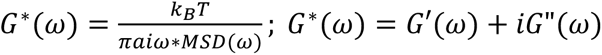, where *ω* is the angular frequency, *MSD*(*ω*) is the Fourier transform of ensemble-time-averaged MSD for each condition, *α* is the probe particle radius. *G*^∗^(*ω*) can be separated into the elastic component as the storage modulus *G*′(*ω*) and the viscous component as the loss modulus *G*′′(*ω*). For simplicity in this study, we chose the temperature *T* to be 37℃ or 310 K. However, to account for ATP-driven non-equilibrium activity in the nucleus, *T* could be replaced by an effective temperature in future work.

To probe long-range active processes in the nucleoplasm and chromatin while minimizing confounds from local viscoelastic heterogeneity, we used two-point microrheology and computed the correlated motion between probe particle pairs (along their line of centers) that separate by a small distance range R^53,107^. For each condition, we averaged over all particles pairs with separation R in each cell and over all cells to obtain the two-point MSD as: 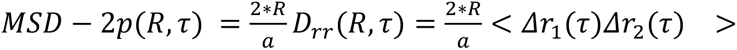, where Δ*r*_1_(*τ*) and Δ*r*_2_(*τ*) are the displacements of the two particles, respectively, along their mutual line of centers in lag time *τ*.

#### Condensate analysis

For analysis of live timelapses of artificial condensates, maximum projections of the z-stack images were pre-processed using FIJI. Condensates were identified and their parameters tracked using the TrackMate plug-in in FIJI^108^. Outputs from TrackMate were filtered and analyzed using a Jupyter notebook in VS Code. For immunofluorescence imaging of fixed condensates, z-stack images were collected with 0.5 μm spacing for a total depth of 20 μm. To generate region of interest (ROI) for marking individual cells, cellpose python package (https://github.com/mouseland/cellpose, ^106^) was used to segment based on the nuclear stain. The resulting images were analyzed with foci-counting, a software package that we are developing in house: https://github.com/liamholtlab/foci_counting.

#### Immunofluorescence

Cells were treated as indicated, then immediately fixed at indicated timepoints with 4% paraformaldehyde (Electron Microscopy Sciences, Cat. No. 15714) for 10 min at room temperature. The cells were subsequently washed three times with 1x PBS, and permeabilized with 0.5% Triton X-100 (Fisher Scientific, Cat. No. 9002-93-1) in 1x PBS for 15 min at room temperature. After blocking with 4% FBS in PBS for 1 h, primary antibodies (1:500 dilution) were applied overnight at 4°C. The next day, the cells were washed three times with 1x PBS and incubated with secondary antibodies (1:500 dilution) in 4% FBS in PBS at room temperature for 1 h in the dark. After washing three times with 1x PBS, cells were stained with SiR-DNA (1:5000 dilution) or 1 μM Hoechst 33342 (Thermo Fisher Scientific 62249). Cells were subsequently stored in 1x PBS at 4°C in the dark until imaging. The fixed plates of cells were mounted on a Nikon spinning disk confocal scanning microscope. Fluorescence signals were obtained using 405 nm, 488 nm, 561 nm, and 640 nm lasers with the following emission filters: 455/50, 525/36, 605/52, and 705/72 (Chroma Technology Corp). Images were captured using a Prime 95B scMOS camera (Photometrics) with a 60x/1.49 numerical aperture objective lens with DIC capability. To calculate cell volume, vRC volume, vRC mean intensity, condensate count, condensate mean intensity, and background intensity, 20-μm Z-stacks were taken with 0.5-μm steps between frames, and images were analyzed with foci-counting.

#### Immunoblotting

Total cellular protein was collected by lysis in sample buffer (62.5 mM Tris-HCl pH 6.8, 2% SDS, 10% glycerol, 0.7M β-mercaptoethanol) followed by boiling for 5 minutes. Lysates were fractionated by sodium dodecyl sulfate polyacrylamide gel electrophoresis (SDS-PAGE) and transferred to nitrocellulose membranes. Membranes were blocked in 5% non-fat milk in TBST for 1 h at room temperature and incubated in primary antibody overnight at 4°C. Primary antibodies were detected using either anti-rabbit IgG HRP or anti-mouse IgG HRP secondary antibodies and visualized by chemiluminescent detection using an Invitrogen iBright FL1000 Imaging System.

## QUANTIFICATION AND STATISTICAL ANALYSIS

Images were quantitated using FIJI plugin MOSAIC for ImageJ^104^. Single particle tracking data were analyzed with GEMspa^105^. Nuclear volumes and vRC volumes were quantified using foci-counting, a software package developed in-house by Sarah Keegan (https://github.com/liamholtlab/foci_counting). Artificial condensates were analyzed and quantified with TrackMate Fiji plug-in^108^. Outputs from TrackMate were filtered and analyzed using a jupyter notebook in VS Code. ChatGPT (GPT-4-turbo, OpenAI) was occasionally used for code debugging and optimization. Graphs were generated by GraphPad Prism 10 (GraphPad Software). Statistical analysis was performed with GraphPad Prism 10 (GraphPad Software). We used the Kruskal-Wallis test for all one-way comparisons. Comparisons with only two groups were performed using a Mann-Whitney test for all data except qPCR data, where a Welch’s t test was utilized. All two-way ANOVAs were followed by Tukey’s post hoc test to correct for multiple comparisons unless otherwise indicated. Further statistical details can be found in the figure legends and in the tables. For all figure legends, n is indicative of the total number of cells analyzed while N is indicative of the number of independent experimental replicates conducted.

## KEY RESOURCES TABLE

| REAGENT | SOURCE | IDENTIFIER |
| --- | --- | --- |
| **Antibodies** |  |  |
| ICP4 | abcam | ab6514 |
| ICP8 | abcam | ab20193 |
| GAPDH | Cell Signaling | 2118S |
| gC | abcam | ab6509 |
| UL42 | Santa Cruz | SC-53331 |
| VP16 | abcam | ab110226 |
| ICP0 | abcam | ab6513 |
| Halo | Promega | G9211 |
| RFP/mScarlet | Rockland | 600-401-379 |
| ICP27 | abcam | ab31631 |
| **Bacterial and Virus Strains** |  |  |
| KOS | Priscilla Schaffer (University of Pennsylvania) | HSV-1 Wild Type |
| 7134 | Priscilla Schaffer (University of Pennsylvania) | HSV-1 ΔICP0 |
| N12 | Neal DeLuca (University of Pittsburgh) | HSV-1 ΔICP4 |
| d22lacZ #8 | Steve Rice (University of Minnesota) | HSV-1 ΔICP22 |
| d27-1 | Steve Rice (University of Minnesota) | HSV-1 ΔICP27 |
| d301 | David Knipe (Harvard Medical School) | HSV-1 ΔICP8 |
| n6 | Neal DeLuca (University of Pittsburgh) | HSV-1 ΔNTA |
| n208 | Neal DeLuca (University of Pittsburgh) | HSV-1 ΔCTA |
| Rosetta (DE3) Competent Cells - Novagen | Millipore Sigma | 70954-3 |
| **Chemicals, Peptides, and Recombinant Proteins** |  |  |
| Cycloheximide | Sigma | C104450 |
| Phosphonoacetic acid, 98% | Sigma-Aldrich | 284270-10G |
| Doxycycline | Sigma-Aldrich | D5207-5G |
| D-Sorbitol | Sigma-Aldrich | S1876 |
| JFX 650 HaloTag Ligand | Janelia/HHMI/Lavis Lab |  |
| TMP-Fluorobenzamide-Halo (TFH) | Huaiying Zhang (Carnegie Mellon University) |  |
| TriPPPPro 2TCOA-dCTP | UHH, AK Meier |  |
| HD653 far red dye | UHH, AK Meier | IK341 |
| Puromycin | MedChem Express | HY-B1743A |
| Polybrene Transfection Reagent | Sigma-Aldrich | TR-1003-G |
| SiR-DNA | Cytoskeleton | CY-SC007 |
| DAPI | Fisher Scientific | D1306 |
| Hoechst 33342 | Thermo Scientific | 62249 |
| G-418 Solution | Sigma-Aldrich | 4727878001 |
| Fugene | Promega | E2312 |
| **Critical Commercial Assays** |  |  |
| Gibson Assembly Master Mix | NEB | E2611S |
| NEBuilder HiFi DNA Assembly Master Mix | NEB | E2621S |
| Q5 DNA Polymerase | NEB | M0491S |
| Phusion DNA Polymerase | Thermo Scientific | F530S |
| **Experimental Models: Cell Lines** |  |  |
| Normal human dermal fibroblasts (NHDF) | Lonza | CC-2509 |
| NHDF-nucPfv (pLH2169 lentivirus) | Liam Holt (NYU Langone Health) | N/A |
| NHDF-50nm-nucGEM (pLH2259 lentivirus) | Liam Holt (NYU Langone Health) | N/A |
| NHDF-Halo-H2A (pLH2498 lentivirus) | Liam Holt (NYU Langone Health) | N/A |
| NHDF-nucPfv-ICP4 (pLH2169 & pLH2488 lentivirus) | Liam Holt (NYU Langone Health) | N/A |
| NHDF-50nm-nucGEM-ICP4 (pLH2259 & pLH2488 lentivirus) | Liam Holt (NYU Langone Health) | N/A |
| NHDF-nucPfv-ICP8 (pLH2169 & pLH2733 lentivirus) | Liam Holt (NYU Langone Health) | N/A |
| NHDF-nucPfv-ICP4-ΔNTA (pLH2169 & pLH2631 lentivirus) | Liam Holt (NYU Langone Health) | N/A |
| NHDF-nucPfv-ICP4-ΔCTA (pLH2169 & pLH2632 lentivirus) | Liam Holt (NYU Langone Health) | N/A |
| NHDF-nucPfv-ICP4-mDBD-RR/LL (pLH2169 & pLH2658 lentivirus) | Liam Holt (NYU Langone Health) | N/A |
| NHDF-nucPfv-ICP4-mDBD-KG/NC (pLH2169 & pLH2659 lentivirus) | Liam Holt (NYU Langone Health) | N/A |
| NHDF-Halo-H2A-ICP4 (pLH2498 & pLH2488 lentivirus) | Liam Holt (NYU Langone Health) | N/A |
| NHDF-Halo-H2A-ICP4-ΔNTA (pLH2498 & pLH2631 lentivirus) | Liam Holt (NYU Langone Health) | N/A |
| HeLa | Jef Boeke (NYU Langone Health) | N/A |
| HeLa-nucPfv (pLH2169 lentivirus) | Liam Holt (NYU Langone Health) | N/A |
| HeLa-50nm-nucGEM (pLH2259 lentivirus) | Liam Holt (NYU Langone Health) | N/A |
| HeLa-12nm-nucGEM (pLH2899 transient transfection) | Liam Holt (NYU Langone Health) | N/A |
| HeLa-nucPfv-ICP4 (pLH2169 lentivirus & pLH2112 transient transfection) | Liam Holt (NYU Langone Health) | N/A |
| HeLa-12nm-nucGEM-ICP4 (pLH2899 & pLH2112 transient transfection) | Liam Holt (NYU Langone Health) | N/A |
| HeLa-ICP4-Mad1-HOTag-miRFP670 (pLH2112, pLH2712, pLH2714 transient transfection) | Liam Holt (NYU Langone Health) | N/A |
| HeLa-ICP4-Mad1-HOTag-eGFP (pLH2112, pLH2713, pLH2714 transient transfection) | Liam Holt (NYU Langone Health) | N/A |
| HEK293T | Jef Boeke (NYU Langone Health) | N/A |
| E5 | Neal DeLuca (University of Pittsburgh) | N/A |
| African green monkey cells (Vero) | ATCC | CCL-81 |
| V22 | Steve Rice (University of Minnesota) | N/A |
| V27 | Steve Rice (University of Minnesota) | N/A |
| V529 | David Knipe (Harvard School of Medicine) | N/A |
| **Recombinant DNA** |  |  |
| pMD2.G | Addgene #12259 | pLH1863 |
| psPAX2 | Addgene #12260 | pLH1864 |
| TetOn-HSV1-ICP4-BFP2 | VectorBuilder custom order: VB211009-1101fac, Addgene #250609 | pLH2112 |
| pUBC-NLS-Pfv-TSapphire | Liam Holt (NYU Langone Health), Addgene # 203653 | pLH2169 |
| pUBC-SV40NLS-popz3h-QtE-GS-TSapphire | Liam Holt (NYU Langone Health), Addgene # 250610 | pLH2259 |
| pUBC-SV40NLS-HoTag3-Shunzhi12nm-Sapphire | Liam Holt (NYU Langone Health), Addgene #250617 | pLH2899 |
| pLVX-TetOn-puro-ICP4 | Liam Holt (NYU Langone Health), Addgene #250611 | pLH2488 |
| pVSV-G | Addgene #138479 | pLH2496 |
| gag/pol | Addgene #14887 | pLH2497 |
| pBABE-puro-Halo-H2A | Eli Rothenberg (NYU Langone Health) | pLH2498 |
| pLVX-TetOn-puro | Meike Dittman (NYU Langone Health) | pLH2537 |
| pLVX-TetOn-puro-ICP4-ΔNTA | Liam Holt (NYU Langone Health),<br>Addgene #250612 | pLH2631 |
| pLVX-TetOne-puro-ICP4-ΔCTA | Liam Holt (NYU Langone Health),<br>Addgene #250613 | pLH2632 |
| pLVX-TetOn-puro-ICP4-mDBD-RR/LL | Liam Holt (NYU Langone Health),<br>Addgene #250614 | pLH2658 |
| pLVX-TetOn-puro-ICP4-mDBD-KG/NC | Liam Holt (NYU Langone Health),<br>Addgene #250615 | pLH2659 |
| p503 jt74 RMCE miRFP670-Halo-HOTag3 | Huaiying Zhang (Carnegie Mellon University) | pLH2712 |
| p403 jt61 RMCE mScarlet-eDHFR-Mad1 | Huaiying Zhang (Carnegie Mellon University) | pLH2714 |
| pLVX-TetOn-puro-ICP8 | Liam Holt (NYU Langone Health),<br>Addgene #250616 | pLH2733 |
| **Software and Algorithms** |  |  |
| Python | Open source | https://www.python.org/downloads/release/python-3107/ |
| FIJI (FIJI is just ImageJ) | 109 | https://imagej.net/software/fiji/ |
| MOSAIC for ImageJ | 104 | https://github.com/mosaic-group/MosaicSuite |
| foci-counting |  | https://github.com/liamholtlab/foci_counting |
| GEMSpa | 105 | https://github.com/liamholtlab/napari-gemspa (napari plugin),<br>https://github.com/liamholtlab/GEMspa (stand alone) |
| Cellpose | 106 | https://cellpose.readthedocs.io/en/latest/# |
| TrackMate | 108 | https://github.com/trackmate-sc/TrackMate |
| Nikon Elements | Nikon Instruments (2017) | https://www.nikoninstruments.com/Products/Software |

## Notes

https://github.com/nlherzog/Biophysical-Tracking

### Competing Interest Statement

The authors have declared no competing interest.

### Summary of Updates

Results section updated to include new two-point microrheology data, complex modulus analysis, and additional tracer particle sizes. Corresponding figures and supplemental files updated (Fig. 1, Fig. 3, Fig. 5, Fig. 7, Fig. S1, Fig. S2, Fig. S3, Fig. S5, Fig. S7, all tables). Authors updated to reflect the addition of these new data and analyses. Discussion updated for clarity.

