## Supplemental Figures and Tables for "Herpes simplex virus 1 fluidizes the nucleus enabling condensate formation"

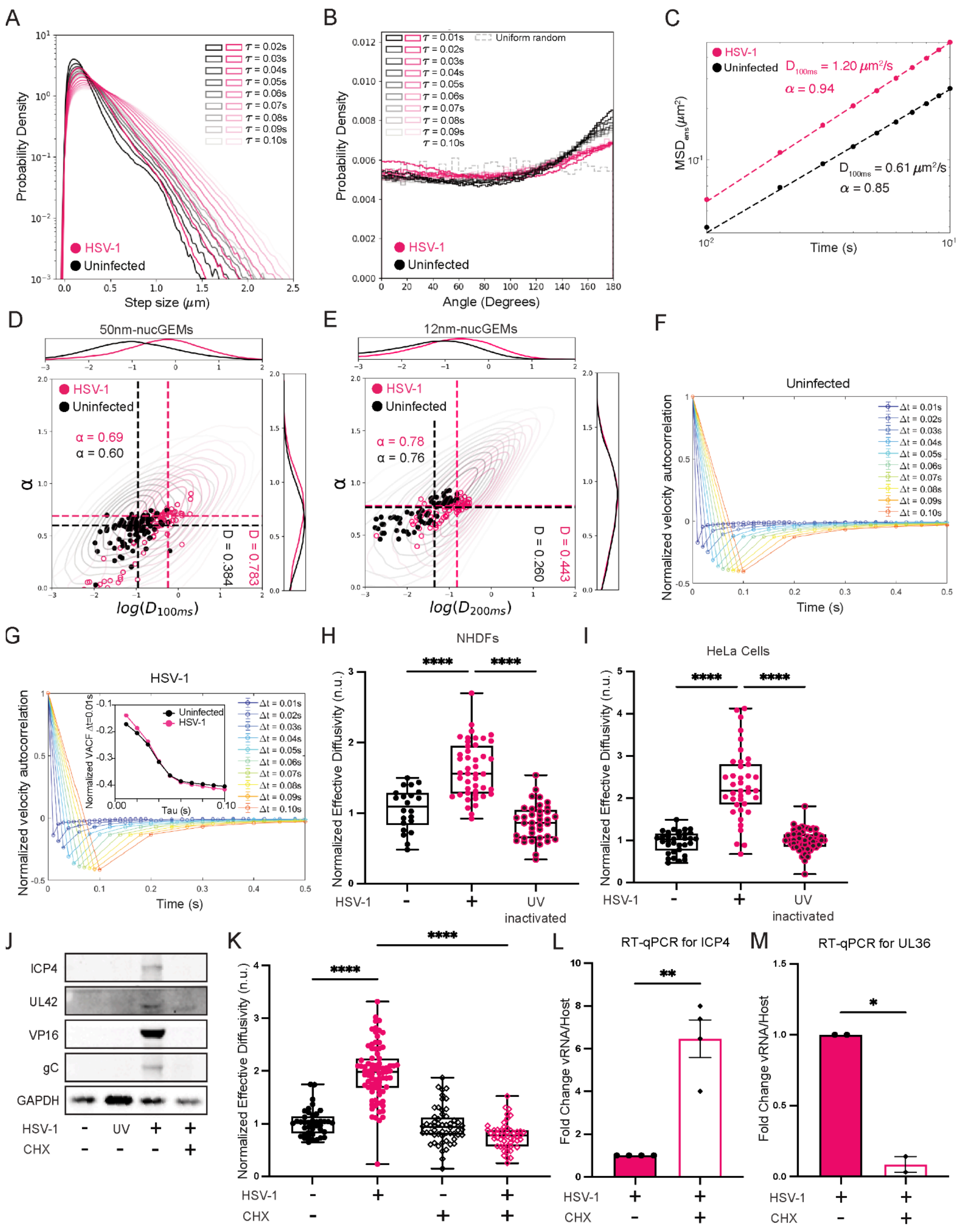
A
B
C
D
E
F
G
H
I
J
K
L
M
HSV-1
Uninfected
Probability Density
Step size (μm)
Angle (Degrees)
Uniform random
D100ms = 1.20 μm²/s
α = 0.94
D100ms = 0.61 μm²/s
α = 0.85
Time (s)
50nm-nucGEMs
12nm-nucGEMs
α = 0.69
α = 0.60
α = 0.78
α = 0.76
D = 0.384
D = 0.783
D = 0.260
D = 0.443
log(D100ms)
log(D200ms)
Normalized velocity autocorrelation
Normalized VACF Δt=0.01s
Tau (s)
NHDFs
HeLa Cells
Normalized Effective Diffusivity (n.u.)
UV inactivated
ICP4
UL42
VP16
gC
GAPDH
CHX
RT-qPCR for ICP4
RT-qPCR for UL36
Fold Change vRNA/Host

**Figure S1: Nuclear diffusivity increases during HSV-1 infection, dependent on new protein synthesis, related to Figure 1.** (A) Step size distribution of 40 nm nucGEM (nucGEM) tracks at different time intervals ($\tau$) in uninfected and HSV-1 infected cells. $\tau$ = 0.01s was excluded due to the presence of artifacts from tracking parameters. n>136; N≥3 biological replicates. (B) Angle distribution of nucGEM tracks from (A) at different time intervals ($\tau$) in uninfected and HSV-1 infected cells. (C) Ensemble-time-averaged mean-squared displacement (MSD) versus time interval ($\tau$) of tracks from (A) on a log-log scale. A linear model was fitted to determine the effective diffusion ($D_{100ms}$) and anomalous exponent (α) values for the indicated conditions. (D,E) Graphical representation of the anomalous exponent (α) compared to the log(effective diffusion) for individual tracks of 50nm-nucGEM (D) and 12nm-nucGEM (E) in HSV-1 infected (pink) or uninfected (dark gray) cells from three representative experiments. Frequency maps along each axis represent the number of cells with a given value. Statistical analysis available in **Table 4**. (F, G) Normalized velocity autocorrelation (VACF) for nucGEM tracks from (A) with respect to $\tau$ = 0. VACFs were calculated from velocities of different time intervals (0-0.1s, blue to red). The inserted graph displays the first negative VACF values, combined from velocities of different time scales for the indicated conditions. (H) Effective diffusivity of nucGEMs in NHDFs infected with WT HSV-1 or UV-inactivated HSV-1 at 9 hpi, MOI =5. n>40; N≥3 biological replicates. (I) Effective diffusivity of nucGEMs in HeLa cells, treated as in (H). n>64; N≥3 biological replicates. (J) Immunoblot against immediate early, early, and late viral proteins in NHDFs 9 hpi with WT HSV-1, UV-inactivated HSV-1 (UV), or HSV-1 with cycloheximide (CHX). Immunoblot is representative of N≥3 biological replicates. (K) Effective diffusivity of nucGEMs in NHDFs treated with CHX and infected with HSV-1 at 9 hpi, MOI = 5. n>82; N≥3 biological replicates. (L) Total RNA was isolated from cells in (K). RT-qPCR analysis was performed for HSV-1 immediate early gene ICP4. n>6; N=3. (M) Total RNA was isolated as in panel L. RT-qPCR analysis was performed for HSV-1 late gene UL36, n>4; N=2. *p<0.05, **p<0.01, ****p<0.0001 by Kruskal-Wallis test (H,I), by two-way ANOVA (K), or by Welch's t test (L,M).

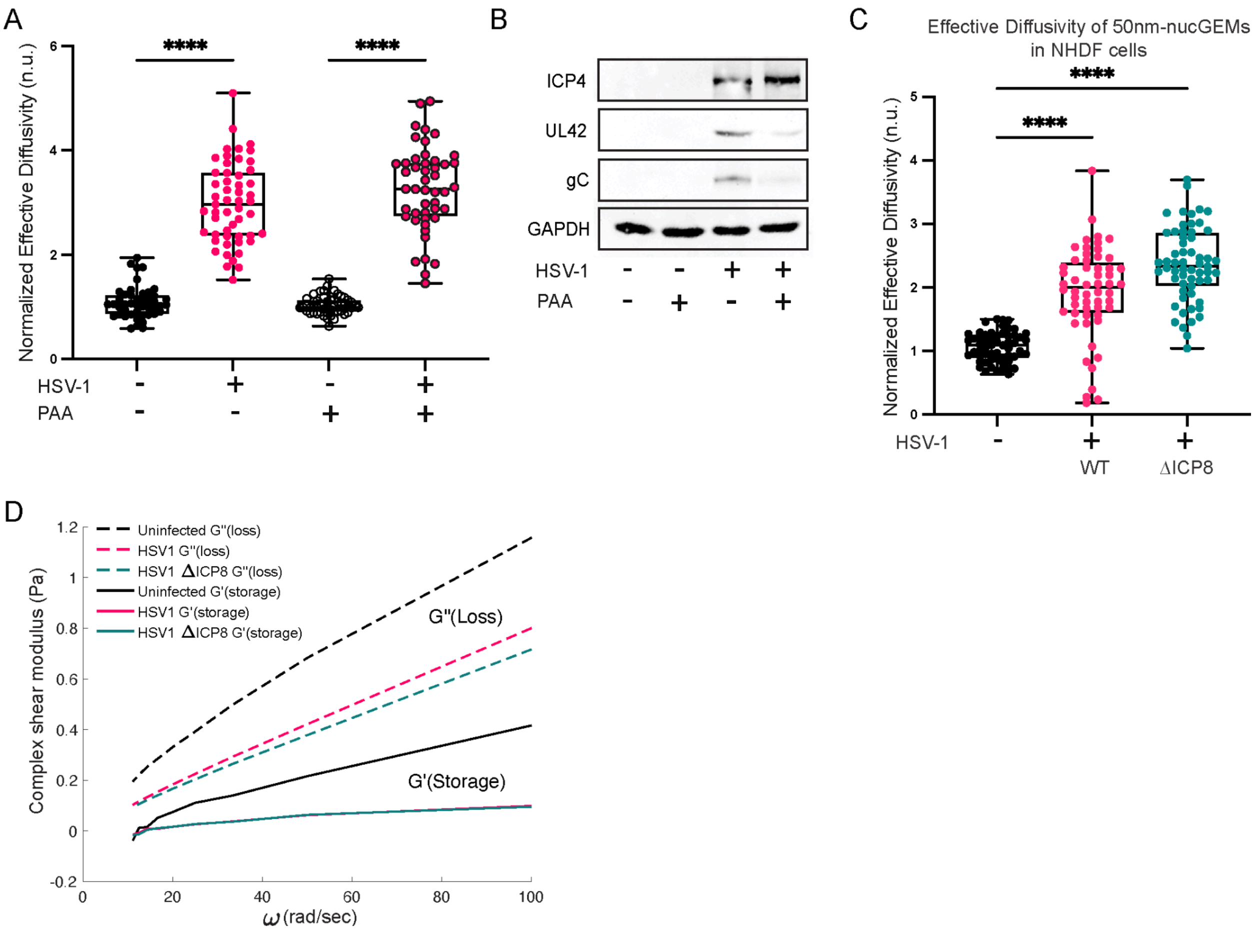
A
Normalized Effective Diffusivity (n.u.)
****
****
HSV-1
PAA
B
ICP4
UL42
gC
GAPDH
HSV-1
PAA
C
Effective Diffusivity of 50nm-nucGEMs in NHDF cells
Normalized Effective Diffusivity (n.u.)
****
****
HSV-1
WT
ΔICP8
D
Uninfected G"(loss)
HSV1 G"(loss)
HSV1 ΔICP8 G"(loss)
Uninfected G'(storage)
HSV1 G'(storage)
HSV1 ΔICP8 G'(storage)
G"(Loss)
G'(Storage)
Complex shear modulus (Pa)
ω (rad/sec)

**Figure S2: HSV-1 infection fluidizes the nucleus independent of viral DNA synthesis, chromatin margination, and nuclear volume changes, related to Figure 2.** (A) HeLa cells expressing nucGEMs were treated with 300 μg/mL phosphonoacetic acid (PAA) and were either left uninfected or infected with WT KOS HSV-1 at an MOI of 5. At 9 hpi, GEM movies and nuclei images were collected and analyzed. n>91; N≥3 biological replicates. (B) Immunoblot analysis to detect ICP4 (immediate-early), UL42 (early), and gC (late) viral protein expression in NHDFs infected for 9 h with WT HSV-1 treated with or without 300 μg/mL PAA. Immunoblot data is representative of N=3 biological replicates. (C) NHDFs expressing 50nm-nucGEMs were infected with WT HSV-1 or HSV-1 ΔICP8 at an MOI of 5. At 9 hpi, GEM movies and nuclei images were obtained and analyzed to determine effective diffusivity as described (Methods). n>139; N≥3 biological replicates. (D) Calculated storage modulus (G', solid lines) and loss modulus (G", dotted lines) in cells from (**Fig. 2A**). ****p<0.0001 by two-way ANOVA (A) or by Kruskal-Wallis test (C).

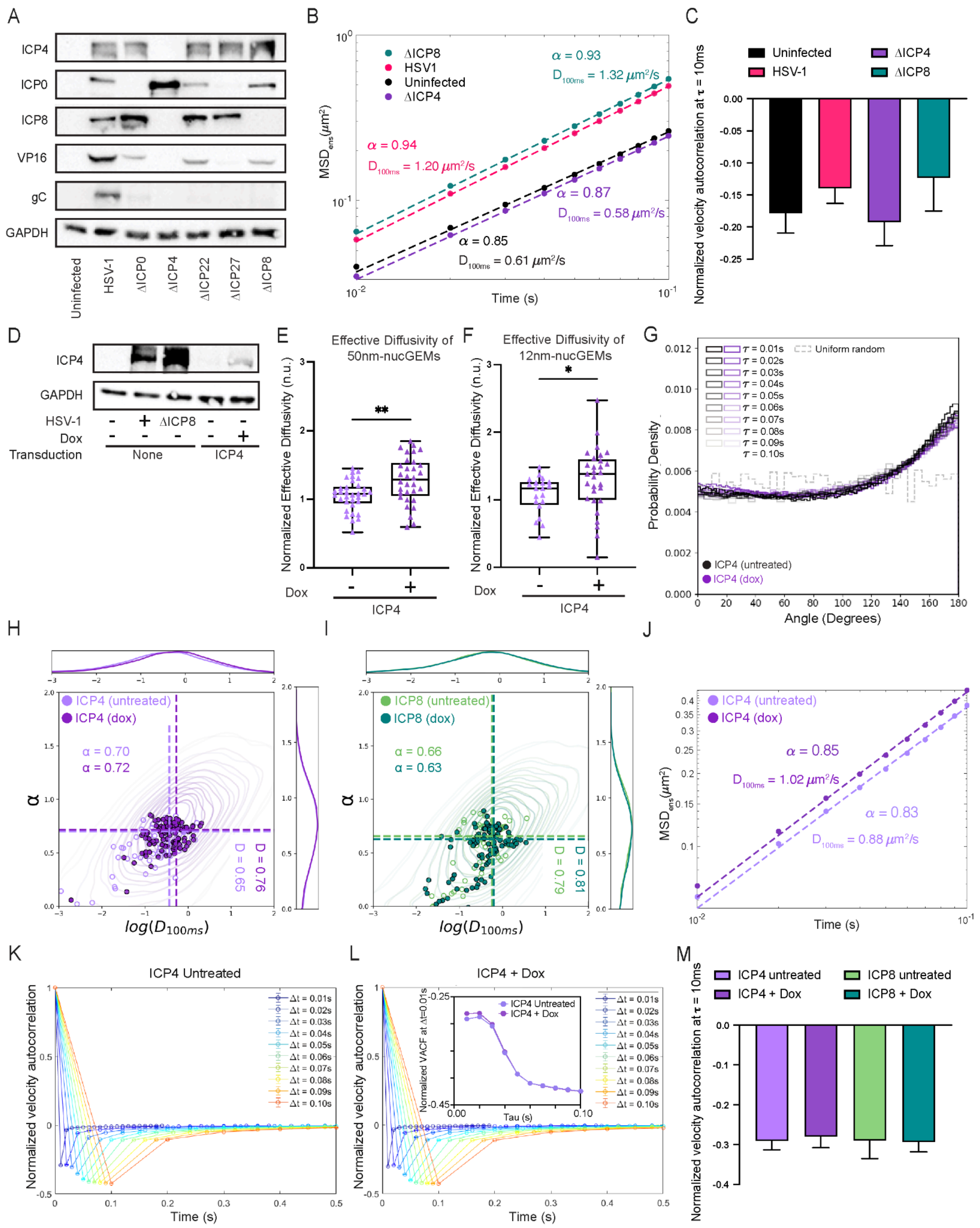

A
ICP4
ICP0
ICP8
VP16
gC
GAPDH
Uninfected
HSV-1
ΔICP0
ΔICP4
ΔICP22
ΔICP27
ΔICP8
B
ΔICP8
HSV1
Uninfected
ΔICP4
α = 0.93
D100ms = 1.32 μm²/s
α = 0.94
D100ms = 1.20 μm²/s
α = 0.87
D100ms = 0.58 μm²/s
α = 0.85
D100ms = 0.61 μm²/s
MSDens (μm²)
Time (s)
C
Uninfected
HSV-1
ΔICP4
ΔICP8
Normalized velocity autocorrelation at τ = 10ms
D
ICP4
GAPDH
HSV-1
ΔICP8
Dox
Transduction
None
ICP4
E
Effective Diffusivity of 50nm-nucGEMs
Normalized Effective Diffusivity (n.u.)
Dox
ICP4
F
Effective Diffusivity of 12nm-nucGEMs
Normalized Effective Diffusivity (n.u.)
Dox
ICP4
G
Uniform random
ICP4 (untreated)
ICP4 (dox)
Probability Density
Angle (Degrees)
H
ICP4 (untreated)
ICP4 (dox)
α = 0.70
α = 0.72
D = 0.76
D = 0.65
log(D100ms)
I
ICP8 (untreated)
ICP8 (dox)
α = 0.66
α = 0.63
D = 0.81
D = 0.79
log(D100ms)
J
ICP4 (untreated)
ICP4 (dox)
α = 0.85
D100ms = 1.02 μm²/s
α = 0.83
D100ms = 0.88 μm²/s
MSDens (μm²)
Time (s)
K
ICP4 Untreated
Normalized velocity autocorrelation
Time (s)
L
ICP4 + Dox
ICP4 Untreated
ICP4 + Dox
Normalized VACF at Δt=0.01s
Tau (s)
Normalized velocity autocorrelation
Time (s)
M
ICP4 untreated
ICP4 + Dox
ICP8 untreated
ICP8 + Dox
Normalized velocity autocorrelation at τ = 10ms

**Figure S3: HSV-1 immediate early protein ICP4 is necessary to fluidize the infected cell nucleus and sufficient to fluidize the nucleus of uninfected cells, related to Figure 3.** (A) NHDFs were infected with WT KOS HSV-1, ΔICP4 HSV-1, ΔICP22 HSV-1, ΔICP27 HSV-1, or ΔICP8 HSV-1 at an MOI of 5, or with ΔICP0 HSV-1 at an MOI of 10. Total protein was collected at 9 hpi, fractionated by SDS-PAGE, and analyzed by immunoblotting against the antibodies shown. GAPDH served as the loading control. Data is representative of N=3 biological replicates. (B) Ensemble-time-averaged mean-squared displacement (MSD) versus time interval ($\tau$) of tracks from (**Fig. 2A**) on a log-log scale. A linear model was fitted to determine the effective diffusion ($D_{100ms}$) and anomalous exponent (α) values for the indicated conditions. (C) Normalized velocity autocorrelation at $\tau$=10 ms 9 hpi with WT HSV-1, ΔICP4 HSV-1, ΔICP8 HSV-1, or uninfected controls. Error bars are standard error of the mean (SEM). (D) NHDFs were transduced as described (Methods) with a lentivirus expressing codon-optimized ICP4 under a tet-promoter or left untransduced. Untransduced cells were then infected with WT HSV-1 KOS or ΔICP8 HSV-1 at an MOI of 5 for 9 h; in transduced cells ICP4 protein expression was induced for 9 h with doxycycline. At 9 hpt total protein was collected and processed as in panel A. Data is representative of N=3 biological replicates. (E) NHDFs expressing 50nm-nucGEMs were transduced as described (Methods) with a lentivirus expressing codon-optimized ICP4 and induced with doxycycline for 9 h. GEM movies were acquired as described (Methods). n>79; N≥3 biological replicates. (F) HeLa cells expressing 12nm-nucGEMs were transfected with a plasmid expressing codon-optimized ICP4 under a tet-promoter and induced with 3 µg/mL doxycycline for 9 h. GEM movies were acquired as described (Methods). n>59; N≥3 biological replicates. (G) Angle distribution of nucGEM tracks from (**Fig. 3E**) at different time intervals ($\tau$) in NHDFs with (Dox) and without (Untreated) ICP4 expression. (H) Graph of anomalous exponent (alpha) compared to the log effective diffusion of tracks in the presence (purple) and absence (grey) of ICP4 induction in NHDFs. Data is pooled from N=4 biological replicates. Frequency maps along each axis represent the number of cells with a given value. (I) Graph as in (H) in the presence (green) and absence (grey) of ICP8 induction in NHDFs. Data is pooled from N=4 biological replicates. Frequency maps along each axis represent the number of cells with a given value. (J) Ensemble-time-averaged mean-squared displacement (MSD) versus time interval ($\tau$) of tracks from (**Fig. 3E**) on a log-log scale, as in (B). (K, L) Normalized velocity autocorrelation (VACF) of tracks from (G, H, J) with respect to $\tau$ = 0 for nucGEMs, with autocorrelations calculated from velocities of different time intervals (0-0.1s, blue to red). The inserted graph displays the first negative VACF values, combined from velocities of different time scales for the indicated conditions. (M) Normalized velocity autocorrelation at $\tau$=10 ms from (**Fig. 3E**). Error bars are standard error of the mean (SEM). Statistical analysis (H, I) available in **Table 5**. *p<0.05, **p<0.01 by Kruskal-Wallis test (C, M) or by Mann-Whitney test (E, F).

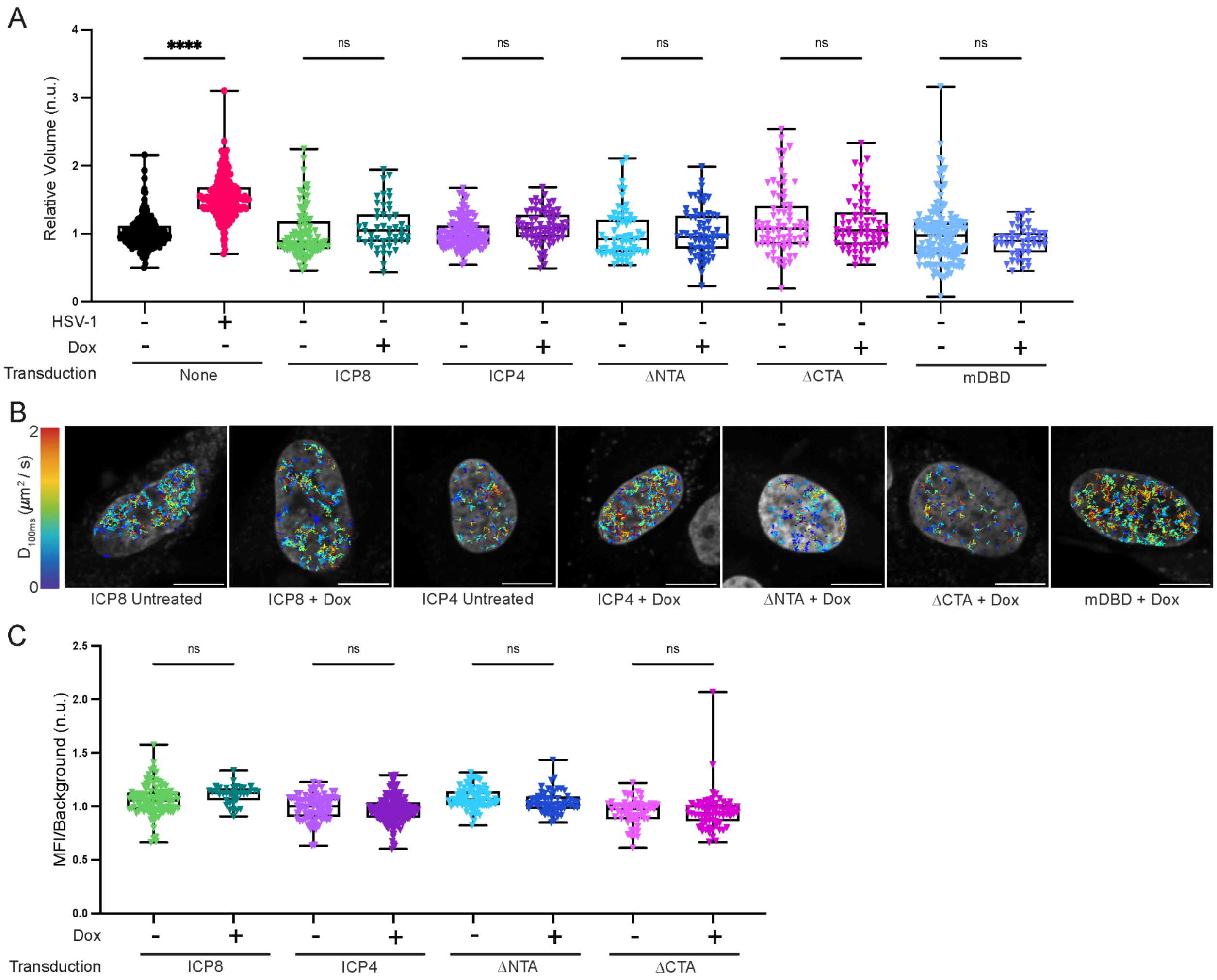
A
Relative Volume (n.u.)
0
1
2
3
4
****
ns
ns
ns
ns
ns
HSV-1
Dox
Transduction
None
ICP8
ICP4
ΔNTA
ΔCTA
mDBD
B
$D_{100ms}$ ($\mu m^2$ / s)
0
2
ICP8 Untreated
ICP8 + Dox
ICP4 Untreated
ICP4 + Dox
ΔNTA + Dox
ΔCTA + Dox
mDBD + Dox
C
MFI/Background (n.u.)
0.0
0.5
1.0
1.5
2.0
2.5
ns
ns
ns
ns
Dox
Transduction
ICP8
ICP4
ΔNTA
ΔCTA

**Figure S4: Both N-terminal and C-terminal domains of ICP4 are required to increase nucGEM diffusivity in uninfected cells, related to Figure 4.** (A) NHDFs expressing nucGEMs were transduced as described (Methods) with lentivirus expressing codon-optimized ICP4, ICP8, or ICP4 mutants described in (**Fig. 4A**) under a tet-promoter. Cells were either infected with WT HSV-1 KOS at an MOI of 5 or protein expression was induced for 9 h. At 9 hpt, cells were stained using vital stain SiR-DNA to visualize the host nucleus. Z-stacks were obtained, nuclear masks were created using cellpose, and relative nuclear volume of cells was calculated with Foci-Counting (Methods). n>90; N≥3 biological replicates. (B) Tracks of individual nucGEMs from representative induced and uninduced NHDFs in conditions as in (A). Tracks are color-coded according to the effective diffusion coefficient. (C) NHDFs were transduced as in panel A. Protein expression was induced for 7 h, then cells were incubated for 2 h with the nucleoside analog 5-ethynyl uridine (5EU). At 9 hpt cells were fixed using PFA and click chemistry was performed. Cells were additionally probed for ICP4 or ICP8 via IF. Cells were verified for ICP4 or ICP8 expression and mean fluorescence intensity (MFI) was calculated using Foci-Counting, then normalized to background (Methods). n>99; N≥3 biological replicates. ****p<0.0001 by Kruskal-Wallis test (all panels).

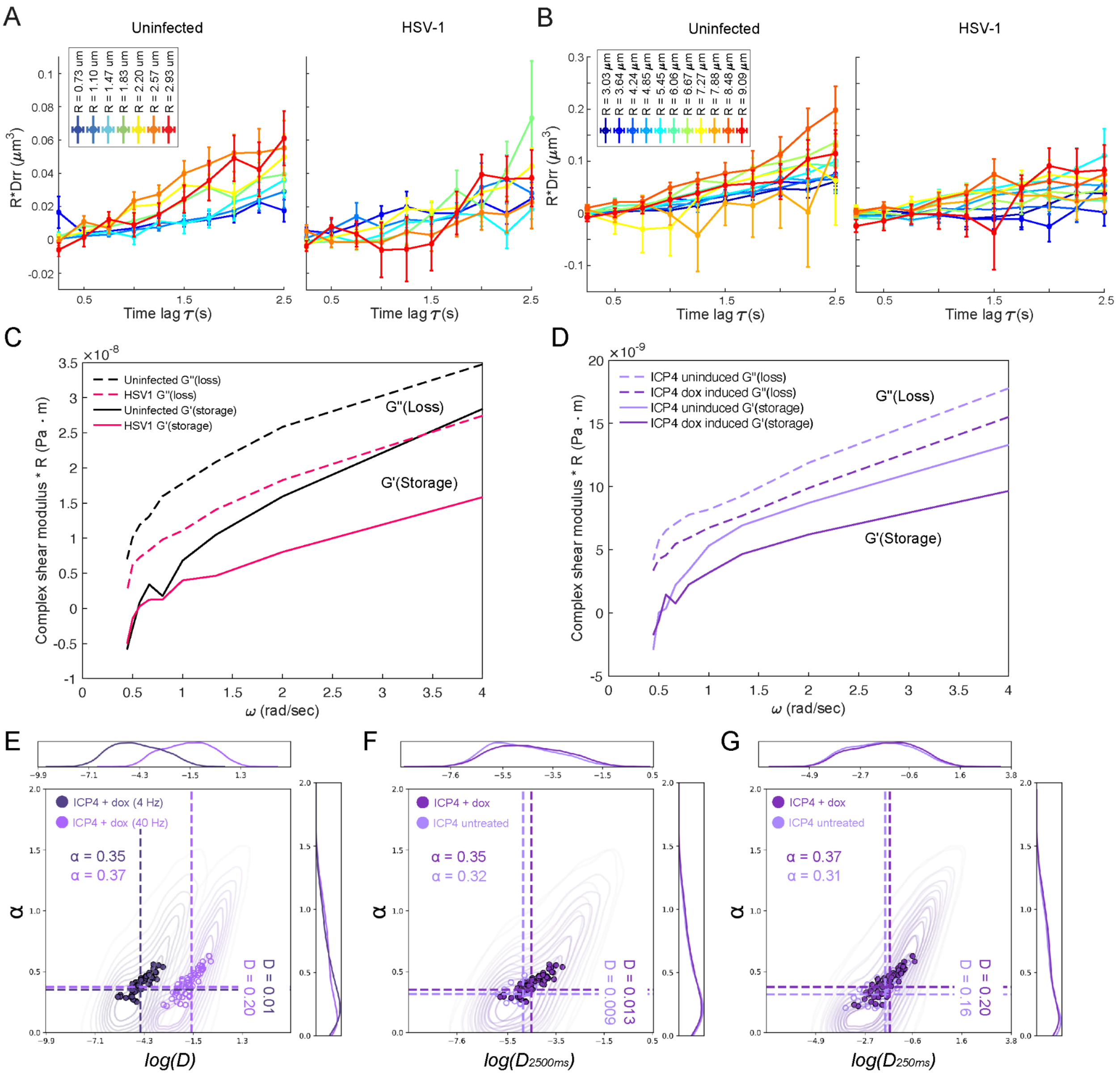
A
Uninfected
HSV-1
R*Drr (μm³)
Time lag τ(s)
R = 0.73 um
R = 1.10 um
R = 1.47 um
R = 1.83 um
R = 2.20 um
R = 2.57 um
R = 2.93 um
B
Uninfected
HSV-1
R*Drr (μm³)
Time lag τ(s)
R = 3.03 μm
R = 3.64 μm
R = 4.24 μm
R = 4.85 μm
R = 5.45 μm
R = 6.06 μm
R = 6.67 μm
R = 7.27 μm
R = 7.88 μm
R = 8.48 μm
R = 9.09 μm
C
×10⁻⁸
Uninfected G"(loss)
HSV1 G"(loss)
Uninfected G'(storage)
HSV1 G'(storage)
G"(Loss)
G'(Storage)
Complex shear modulus * R (Pa · m)
ω (rad/sec)
D
×10⁻⁹
ICP4 uninduced G"(loss)
ICP4 dox induced G"(loss)
ICP4 uninduced G'(storage)
ICP4 dox induced G'(storage)
G"(Loss)
G'(Storage)
Complex shear modulus * R (Pa · m)
ω (rad/sec)
E
ICP4 + dox (4 Hz)
ICP4 + dox (40 Hz)
α = 0.35
α = 0.37
D = 0.01
D = 0.20
α
log(D)
F
ICP4 + dox
ICP4 untreated
α = 0.35
α = 0.32
D = 0.013
D = 0.009
α
log(D2500ms)
G
ICP4 + dox
ICP4 untreated
α = 0.37
α = 0.31
D = 0.20
D = 0.16
α
log(D250ms)

**Figure S5: Movement of bound histones increases during ICP4 induction and during HSV-1 infection dependent on the presence of ICP4, related to Figure 5.** (A) Two-point microrheology (TPM) as shown previously quantifying the correlated motion for particle pairs separated by a distance range [R] in uninfected and HSV-1 infected NHDFs identified in **Fig. 5C** (B) TPM as in (A) with an increased distance range [R]. (C) Calculated storage modulus (G', solid lines) and loss modulus (G", dotted lines) from histone tracking in **Fig. 5C.** (D) Calculated storage modulus (G', solid lines) and loss modulus (G", dotted lines) from histone tracking in **Fig. 5E.** (E) Graph of anomalous exponent (alpha) compared to the log effective diffusion of Halo-H2A tracks at 4 Hz (purple) and 40 Hz (grey) of ICP4 induction in NHDFs. (F) Graph of anomalous exponent (alpha) compared to the log effective diffusion of Halo-H2A tracks at 4 Hz in the presence (purple) and absence (grey) of ICP4 induction in NHDFs. (G) Graph of anomalous exponent (alpha) compared to the log effective diffusion of Halo-H2A tracks at 40 Hz in the presence (purple) and absence (grey) of ICP4 induction in NHDFs. Frequency maps along each axis represent the number of cells with a given value. All data is pooled from N=4 biological replicates. Statistical analysis (E, F, G) available in **Table 6**.

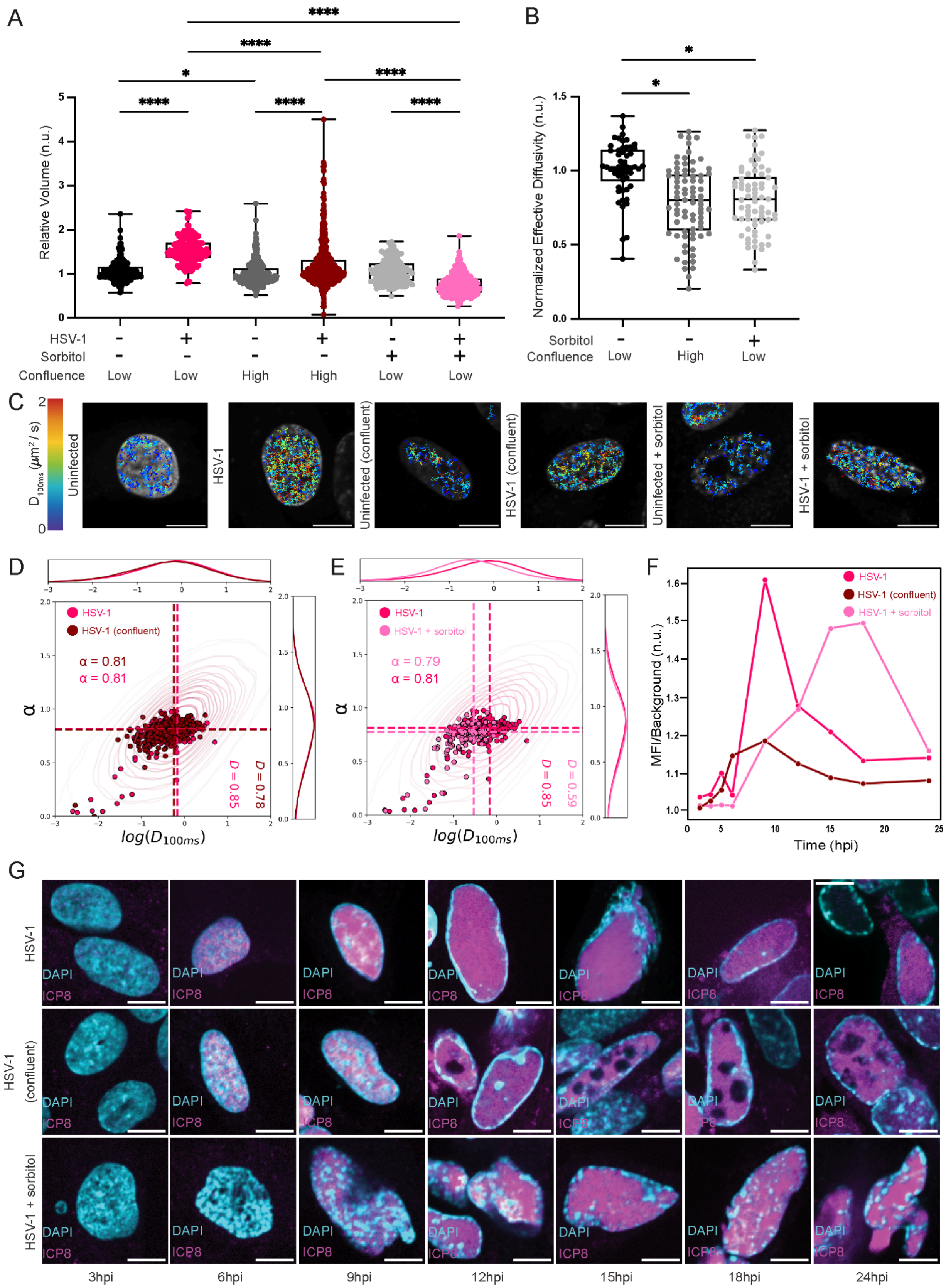
A
B
C
D
E
F
G
Relative Volume (n.u.)
HSV-1
Sorbitol
Confluence
Low
High
Normalized Effective Diffusivity (n.u.)
Uninfected
HSV-1
Uninfected (confluent)
HSV-1 (confluent)
Uninfected + sorbitol
HSV-1 + sorbitol
HSV-1 (confluent)
HSV-1 + sorbitol
α = 0.81
α = 0.81
α = 0.79
α = 0.81
D = 0.85
D = 0.78
D = 0.85
D = 0.59
log(D100ms)
α
MFI/Background (n.u.)
Time (hpi)
DAPI
ICP8
3hpi
6hpi
9hpi
12hpi
15hpi
18hpi
24hpi

**Figure S6: Preventing nuclear fluidization disrupts viral replication compartment formation and decreases infectious virus production, related to Figure 6.** (A) NHDFs transduced with nucGEMs were grown to a confluent monolayer (High), then infected with WT HSV-1 at MOI = 5. NHDFs transduced with nucGEMs were grown to 70% confluency, then infected with WT HSV-1 at MOI = 5 (Low). At 2.5 hpi, cells were left untreated or were treated with 150 mM sorbitol. At 9 hpi cells were stained using vital stain SiR-DNA to visualize the host nucleus. Z-stacks were obtained, nuclear masks were created using cellpose, and relative nuclear volume of cells was calculated with Foci-Counting (Methods). $n>353$; $N\geq3$ biological replicates. (B) NHDFs transduced with nucGEMs were grown to a confluent monolayer (High) or were grown to 70% confluency (Low) and either left untreated or were treated with 150 mM sorbitol. nucGEM diffusivity was measured at 6.5 h as previously described (Methods). $n>136$; $N\geq3$ biological replicates. (C)Tracks of individual nucGEMs from representative infected and uninfected NHDFs under each treatment condition, color-coded according to the effective diffusion coefficient. (D) Graph of anomalous exponent (alpha) compared to the log effective diffusion of tracks during WT HSV-1 infection (HSV-1) vs. WT HSV-1 infection in confluent monolayers (HSV-1 (confluent)). Data is pooled from N=4 biological replicates. Frequency maps along each axis represent the number of cells with a given value. (E) Graph of anomalous exponent (alpha) compared to the log effective diffusion of tracks during WT HSV-1 infection (HSV-1) vs. WT HSV-1 infection with sorbitol treatment as described in (A) (HSV-1 + sorbitol). Data is pooled from N=4 biological replicates. Frequency maps along each axis represent the number of cells with a given value. (F) Mean fluorescence intensity (MFI) of ICP8 indirect immunofluorescence detection within each nucleus in (**Fig. 6D**) was calculated using Foci-Counting and normalized to background. Median MFI across all replicates was plotted as a stacked line graph. (G) Representative immunofluorescence from cells in panel F. Statistical analysis (D, E) available in **Table 7**. All experiments are at least N=3 biological replicates. IF images are representative of N=3 biological replicates. *$p<0.05$, ****$p<0.0001$ by two-way ANOVA with Bonferroni's correction (A) or by Kruskal-Wallis test (B).

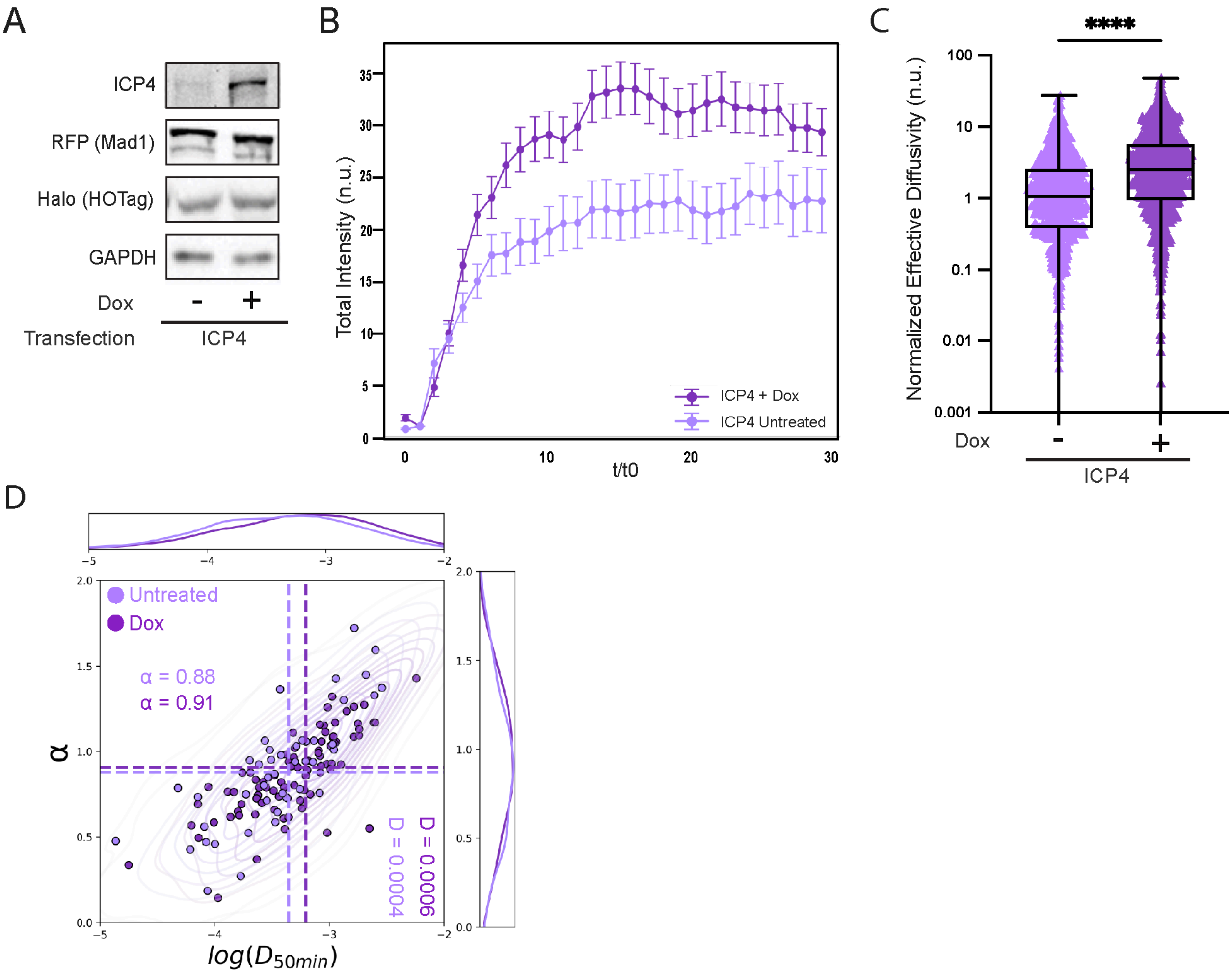
A
ICP4
RFP (Mad1)
Halo (HOTag)
GAPDH
Dox
-
+
Transfection
ICP4
B
Total Intensity (n.u.)
t/t0
ICP4 + Dox
ICP4 Untreated
C
****
Normalized Effective Diffusivity (n.u.)
100
10
1
0.1
0.01
0.001
Dox
-
+
ICP4
D
Untreated
Dox
α = 0.88
α = 0.91
D = 0.0006
D = 0.0004
α
$log(D_{50min})$

**Figure S7: Fluidization of the nucleus by ICP4 facilitates growth of artificial condensates, related to Figure 7.** (A) Immunoblot against ICP4 and the components of the artificial condensate system as described (**Fig. 7A**, Methods). HeLa cells were transfected with the plasmid containing inducible codon-optimized ICP4 as well as the two plasmids forming the artificial condensate system as described (Methods). Cells were treated with 3 μg/mL doxycycline for 8 h or left untreated. At 8 hpt, all conditions were treated with 50 nM chemical dimerizer Trimethoprim-Fluorobenzamide-Halo ligand (TFH) for 1 h. After 1 h of TFH treatment, total protein was collected, fractionated by SDS-PAGE, and analyzed by immunoblotting using the antibodies shown. GAPDH was used as a loading control. Data is representative of N=3 biological replicates. (B) Quantitation of average total intensity of each condensate over time for condensates in cells with (+Dox) or without (untreated) ICP4 expression. Cells were prepared and treated as in panel A. Total condensate intensity for each individual condensate was normalized to the mean intensity of the nucleus in the diffuse phase (pre-TFH treatment), and time was normalized to the timepoint at which condensates nucleated (t/t0). Statistical analysis for (B) available in **Table 8**. (C) Cells were treated as in (A), then imaged every 5 minutes after TFH treatment for 3 h. Condensates were identified and binned according to their mean fluorescence intensity. The bin containing the highest number of condensates with a track at least 10 steps in length was then analyzed for effective diffusivity as previously described (Methods). (D) Graph of anomalous exponent (alpha) compared to the log(effective diffusion) for individual tracks of artificial condensates in the presence (dark purple) and absence (light purple) of ICP4 induction in HeLa cells. Statistical analysis for (D) available in **Table 9**. ****p<0.0001 by Mann-Whitney test (C,D).

**Table 1: Welch's t-test of Log($D_{100ms}$) and α, related to Figure 1G**

| Welch's t-test of Log($D_{100ms}$) and α (alpha) | | | | | | | |
|---|---|---|---|---|---|---|---|
| **variable** | **Group 1** | **Group2** | **Mean group1** | **Mean group2** | **t_statistic** | **p_value** | **significant** |
| **log_D** | Uninfected | HSV-1 | -0.818024253 | -0.129086091 | -141.1539303 | p<0.0001 | TRUE |
| **alpha** | Uninfected | HSV-1 | 0.692879712 | 0.796449879 | -54.85629231 | p<0.0001 | TRUE |

**Table 2: Two-way ANOVA of vRC CTCF, related to Figure 6D**

| Two-way ANOVA of vRC CTCF | | | | | | |
|---|---|---|---|---|---|---|
| **ANOVA table** | **SS (Type III)** | **DF** | **MS** | **F (DFn, DFd)** | | **P value** |
| **Row Factor** | 1.71275E+13 | 591 | 28980620449 | F (591, 15870) = 0.8522 | | 0.9956 |
| **Column Factor** | 1.337E+14 | 26 | 5.14201E+12 | F (26, 15870) = 151.2 | | <0.0001 |
| **Residual** | 5.397E+14 | 15870 | 34007755147 | | | |
| **Multiple Comparison of Mean vRC CTCF — Tukey HSD, FWER=0.05** | | | | | | |
| **Group1** | **Group2** | **MeanDiff** | **p-adj** | **lower** | **upper** | **reject** |
| HSV-1 3hpi | HSV-1 4hpi | -16716 | >0.9999 | -67401 | 33969 | FALSE |
| HSV-1 3hpi | HSV-1 5hpi | -47772 | 0.0672 | -96733 | 1189 | FALSE |
| HSV-1 3hpi | HSV-1 6hpi | -160181 | <0.0001 | -204622 | -115741 | TRUE |
| HSV-1 3hpi | HSV-1 9hpi | -360779 | <0.0001 | -404599 | -316958 | TRUE |
| HSV-1 3hpi | HSV-1 12hpi | -150462 | <0.0001 | -199799 | -101124 | TRUE |
| HSV-1 3hpi | HSV-1 15hpi | -115509 | <0.0001 | -162303 | -68716 | TRUE |
| HSV-1 3hpi | HSV-1 18hpi | -65899 | 0.0007 | -117449 | -14350 | TRUE |
| HSV-1 3hpi | HSV-1 24hpi | -101901 | <0.0001 | -151869 | -51933 | TRUE |
| HSV-1 + sorbitol 3hpi | HSV-1 + sorbitol 4hpi | -1098 | >0.9999 | -53036 | 50840 | FALSE |
| HSV-1 + sorbitol 3hpi | HSV-1 + sorbitol 5hpi | -697 | >0.9999 | -52220 | 50826 | FALSE |
| HSV-1 + sorbitol 3hpi | HSV-1 + sorbitol 6hpi | -60452 | 0.0003 | -105706 | -15198 | TRUE |
| HSV-1 + sorbitol 3hpi | HSV-1 + sorbitol 9hpi | -170370 | <0.0001 | -214316 | -126425 | TRUE |
| HSV-1 + sorbitol 3hpi | HSV-1 + sorbitol 12hpi | -145241 | <0.0001 | -195965 | -94516 | TRUE |
| HSV-1 + sorbitol 3hpi | HSV-1 + sorbitol 15hpi | -209909 | <0.0001 | -258959 | -160860 | TRUE |
| HSV-1 + sorbitol 3hpi | HSV-1 + sorbitol 18hpi | -254410 | <0.0001 | -306035 | -202785 | TRUE |
| HSV-1 + sorbitol 3hpi | HSV-1 + sorbitol 24hpi | -373007 | <0.0001 | -427816 | -318197 | TRUE |
| HSV-1 (confluent) 3hpi | HSV-1 (confluent) 4hpi | -12647 | >0.9999 | -58128 | 32833 | FALSE |
| HSV-1 (confluent) 3hpi | HSV-1 (confluent) 5hpi | -27358 | 0.8943 | -72708 | 17991 | FALSE |
| HSV-1 (confluent) 3hpi | HSV-1 (confluent) 6hpi | -152149 | <0.0001 | -192730 | -111569 | TRUE |
| HSV-1 (confluent) 3hpi | HSV-1 (confluent) 9hpi | -167023 | <0.0001 | -205494 | -128552 | TRUE |
| HSV-1 (confluent) 3hpi | HSV-1 (confluent) 12hpi | -117970 | <0.0001 | -159046 | -76893 | TRUE |
| HSV-1 (confluent) 3hpi | HSV-1 (confluent) 15hpi | -102890 | <0.0001 | -144418 | -61362 | TRUE |
| HSV-1 (confluent) 3hpi | HSV-1 (confluent) 18hpi | -91494 | <0.0001 | -133224 | -49763 | TRUE |
| HSV-1 (confluent) 3hpi | HSV-1 (confluent) 24hpi | -86807 | <0.0001 | -128137 | -45476 | TRUE |

| | | | | | | |
|---|---|---|---|---|---|---|
| HSV-1 3hpi | HSV-1 + sorbitol 3hpi | 20213 | 0.9998 | -32121 | 72548 | FALSE |
| HSV-1 3hpi | HSV-1 (confluent) 3hpi | 15647 | >0.9999 | -33763 | 65057 | FALSE |
| HSV-1 4hpi | HSV-1 + sorbitol 4hpi | 35831 | 0.6267 | -14401 | 86064 | FALSE |
| HSV-1 4hpi | HSV-1 (confluent) 4hpi | 19716 | 0.9991 | -27429 | 66861 | FALSE |
| HSV-1 5hpi | HSV-1 + sorbitol 5hpi | 67289 | <0.0001 | 19191 | 115386 | TRUE |
| HSV-1 5hpi | HSV-1 (confluent) 5hpi | 36061 | 0.3686 | -8977 | 81099 | FALSE |
| HSV-1 6hpi | HSV-1 + sorbitol 6hpi | 119943 | <0.0001 | 84534 | 155352 | TRUE |
| HSV-1 6hpi | HSV-1 (confluent) 6hpi | 23679 | 0.6919 | -10588 | 57946 | FALSE |
| HSV-1 9hpi | HSV-1 + sorbitol 9hpi | 210622 | <0.0001 | 177686 | 243557 | TRUE |
| HSV-1 9hpi | HSV-1 (confluent) 9hpi | 209403 | <0.0001 | 178559 | 240247 | TRUE |
| HSV-1 12hpi | HSV-1 + sorbitol 12hpi | 25434 | 0.9695 | -21951 | 72820 | FALSE |
| HSV-1 12hpi | HSV-1 (confluent) 12hpi | 48140 | 0.0042 | 7144 | 89135 | TRUE |
| HSV-1 15hpi | HSV-1 + sorbitol 15hpi | -74187 | <0.0001 | -116719 | -31654 | TRUE |
| HSV-1 15hpi | HSV-1 (confluent) 15hpi | 28267 | 0.5281 | -9631 | 66165 | FALSE |
| HSV-1 18hpi | HSV-1 + sorbitol 18hpi | -168297 | <0.0001 | -218987 | -117608 | TRUE |
| HSV-1 18hpi | HSV-1 (confluent) 18hpi | -9947 | >0.9999 | -54322 | 34428 | FALSE |
| HSV-1 24hpi | HSV-1 + sorbitol 24hpi | -250892 | <0.0001 | -303496 | -198289 | TRUE |
| HSV-1 24hpi | HSV-1 (confluent) 24hpi | 30741 | 0.5569 | -11005 | 72488 | FALSE |

**Table 3: Independent t-test of condensate count and R/R0, related to Figure 7C-D**

| Independent t-test of condensate count per cell | | | | | | | |
|---|---|---|---|---|---|---|---|
| **Group 1** | **Group 2** | **t/t0** | **Group 1 Mean** | **Group 2 Mean** | **t-statistic** | **p-value** | **Reject** |
| ICP4 + Dox | ICP4 Untreated | 0 | 2.352941176 | 1.88 | 1.168882456 | 0.247796811 | FALSE |
| ICP4 + Dox | ICP4 Untreated | 1 | 3.189189189 | 2.62962963 | 0.992582707 | 0.325111173 | FALSE |
| ICP4 + Dox | ICP4 Untreated | 2 | 4.4 | 4.419354839 | -0.022541243 | 0.982090981 | FALSE |
| ICP4 + Dox | ICP4 Untreated | 3 | 4.960784314 | 6.578947368 | -1.553284158 | 0.125219085 | FALSE |
| ICP4 + Dox | ICP4 Untreated | 4 | 5.866666667 | 7.234042553 | -1.38395612 | 0.169720645 | FALSE |
| ICP4 + Dox | ICP4 Untreated | 5 | 7.030769231 | 8.019230769 | -1.021024169 | 0.309651037 | FALSE |
| ICP4 + Dox | ICP4 Untreated | 6 | 8.060606061 | 8.298245614 | -0.246283946 | 0.805904149 | FALSE |
| ICP4 + Dox | ICP4 Untreated | 7 | 8.909090909 | 8.694915254 | 0.220407351 | 0.825934144 | FALSE |
| ICP4 + Dox | ICP4 Untreated | 8 | 8.956521739 | 8.868852459 | 0.090373431 | 0.928136641 | FALSE |
| ICP4 + Dox | ICP4 Untreated | 9 | 9.242857143 | 9.161290323 | 0.084227291 | 0.933009573 | FALSE |
| ICP4 + Dox | ICP4 Untreated | 10 | 9.464788732 | 9.285714286 | 0.182816459 | 0.855230248 | FALSE |
| ICP4 + Dox | ICP4 Untreated | 11 | 9.662162162 | 9.555555556 | 0.107943636 | 0.914206165 | FALSE |
| ICP4 + Dox | ICP4 Untreated | 12 | 9.723684211 | 9.857142857 | -0.135674962 | 0.892285912 | FALSE |
| ICP4 + Dox | ICP4 Untreated | 13 | 9.658227848 | 10.23809524 | -0.595260444 | 0.552676108 | FALSE |
| ICP4 + Dox | ICP4 Untreated | 14 | 9.962025316 | 10.47619048 | -0.514700004 | 0.60760846 | FALSE |
| ICP4 + Dox | ICP4 Untreated | 15 | 10.16049383 | 10.71428571 | -0.548998339 | 0.58391125 | FALSE |
| ICP4 + Dox | ICP4 Untreated | 16 | 10.37037037 | 10.67692308 | -0.295592386 | 0.767989725 | FALSE |
| ICP4 + Dox | ICP4 Untreated | 17 | 10.63855422 | 10.90769231 | -0.257107702 | 0.797481831 | FALSE |
| ICP4 + Dox | ICP4 Untreated | 18 | 11 | 11.12121212 | -0.111104884 | 0.911694745 | FALSE |
| ICP4 + Dox | ICP4 Untreated | 19 | 11.32142857 | 11.43939394 | -0.105874899 | 0.915834159 | FALSE |
| ICP4 + Dox | ICP4 Untreated | 20 | 11.6 | 11.75757576 | -0.141410222 | 0.887747277 | FALSE |
| ICP4 + Dox | ICP4 Untreated | 21 | 11.98823529 | 11.93939394 | 0.042904221 | 0.96583827 | FALSE |
| ICP4 + Dox | ICP4 Untreated | 22 | 12.43529412 | 12.24242424 | 0.163981623 | 0.869973873 | FALSE |
| ICP4 + Dox | ICP4 Untreated | 23 | 12.64705882 | 12.68181818 | -0.029187114 | 0.976755656 | FALSE |
| ICP4 + Dox | ICP4 Untreated | 24 | 13.05882353 | 13.01515152 | 0.035728697 | 0.971548403 | FALSE |
| ICP4 + Dox | ICP4 Untreated | 25 | 13.50588235 | 13.34848485 | 0.12550164 | 0.900302986 | FALSE |
| ICP4 + Dox | ICP4 Untreated | 26 | 13.98823529 | 13.57575758 | 0.317165738 | 0.751580328 | FALSE |
| ICP4 + Dox | ICP4 Untreated | 27 | 14.28235294 | 13.84848485 | 0.32507335 | 0.745604611 | FALSE |
| ICP4 + Dox | ICP4 Untreated | 28 | 14.28235294 | 13.84848485 | 0.32507335 | 0.745604611 | FALSE |
| ICP4 + Dox | ICP4 Untreated | 29 | 14.28235294 | 13.84848485 | 0.32507335 | 0.745604611 | FALSE |
| ICP4 + Dox | ICP4 Untreated | 30 | 14.28235294 | 13.84848485 | 0.32507335 | 0.745604611 | FALSE |
| **Independent t-test of R/R0** | | | | | | | |
| **Group 1** | **Group 2** | **t/t0** | **Group 1 Mean** | **Group 2 Mean** | **t-statistic** | **p-value** | **Reject** |
| ICP4 + Dox | ICP4 Untreated | 0 | 1.174967089 | 0.778012096 | 5.198373393 | 5.24E-07 | TRUE |
| ICP4 + Dox | ICP4 Untreated | 1 | 1 | 1 | - | - | - |
| ICP4 + Dox | ICP4 Untreated | 2 | 1.537614063 | 1.863523798 | -1.97756027 | 0.048969876 | TRUE |
| ICP4 + Dox | ICP4 Untreated | 3 | 2.230352467 | 2.16533689 | 0.359929475 | 0.719144548 | FALSE |
| ICP4 + Dox | ICP4 Untreated | 4 | 2.98419341 | 2.480592537 | 2.667519572 | 0.008042592 | TRUE |
| ICP4 + Dox | ICP4 Untreated | 5 | 3.31378591 | 2.650254102 | 3.274413685 | 0.001178208 | TRUE |

| | | | | | | | |
|---|---|---|---|---|---|---|---|
| ICP4 + Dox | ICP4 Untreated | 6 | 3.446338921 | 2.7887401 | 3.193641354 | 0.001552965 | TRUE |
| ICP4 + Dox | ICP4 Untreated | 7 | 3.662189308 | 2.767026316 | 4.348247686 | 1.87E-05 | TRUE |
| ICP4 + Dox | ICP4 Untreated | 8 | 3.740582452 | 2.78025408 | 4.584545131 | 6.67E-06 | TRUE |
| ICP4 + Dox | ICP4 Untreated | 9 | 3.806296834 | 2.789549812 | 5.000343226 | 9.68E-07 | TRUE |
| ICP4 + Dox | ICP4 Untreated | 10 | 3.789500958 | 2.832477584 | 4.566930928 | 7.23E-06 | TRUE |
| ICP4 + Dox | ICP4 Untreated | 11 | 3.74619641 | 2.822884107 | 4.448149286 | 1.23E-05 | TRUE |
| ICP4 + Dox | ICP4 Untreated | 12 | 3.789499984 | 2.841257303 | 4.561422549 | 7.40E-06 | TRUE |
| ICP4 + Dox | ICP4 Untreated | 13 | 3.927816501 | 2.866956381 | 4.843163236 | 2.05E-06 | TRUE |
| ICP4 + Dox | ICP4 Untreated | 14 | 3.912022442 | 2.856061222 | 4.790705411 | 2.62E-06 | TRUE |
| ICP4 + Dox | ICP4 Untreated | 15 | 3.920897003 | 2.83438289 | 5.066593989 | 7.07E-07 | TRUE |
| ICP4 + Dox | ICP4 Untreated | 16 | 3.932064087 | 2.840346632 | 5.082926439 | 6.55E-07 | TRUE |
| ICP4 + Dox | ICP4 Untreated | 17 | 3.836400105 | 2.860668019 | 4.503137618 | 9.58E-06 | TRUE |
| ICP4 + Dox | ICP4 Untreated | 18 | 3.778740712 | 2.858336512 | 4.322670721 | 2.10E-05 | TRUE |
| ICP4 + Dox | ICP4 Untreated | 19 | 3.75108435 | 2.870119005 | 4.077457992 | 5.88E-05 | TRUE |
| ICP4 + Dox | ICP4 Untreated | 20 | 3.75185213 | 2.82494463 | 4.371839811 | 1.70E-05 | TRUE |
| ICP4 + Dox | ICP4 Untreated | 21 | 3.767321397 | 2.776276119 | 4.656496913 | 4.82E-06 | TRUE |
| ICP4 + Dox | ICP4 Untreated | 22 | 3.808600949 | 2.820064372 | 4.679783807 | 4.35E-06 | TRUE |
| ICP4 + Dox | ICP4 Untreated | 23 | 3.77975886 | 2.808503929 | 4.612313841 | 5.91E-06 | TRUE |
| ICP4 + Dox | ICP4 Untreated | 24 | 3.785988717 | 2.850876804 | 4.357800157 | 1.82E-05 | TRUE |
| ICP4 + Dox | ICP4 Untreated | 25 | 3.76613203 | 2.844883132 | 4.29399812 | 2.38E-05 | TRUE |
| ICP4 + Dox | ICP4 Untreated | 26 | 3.779103641 | 2.873360563 | 4.146192261 | 4.43E-05 | TRUE |
| ICP4 + Dox | ICP4 Untreated | 27 | 3.686014666 | 2.778421836 | 4.37230407 | 1.71E-05 | TRUE |
| ICP4 + Dox | ICP4 Untreated | 28 | 3.63253516 | 2.786234598 | 3.909482604 | 0.000115181 | TRUE |
| ICP4 + Dox | ICP4 Untreated | 29 | 3.608946765 | 2.783536085 | 3.874770058 | 0.000132257 | TRUE |

**Table 4: Welch's t-test of Log($D_{100ms}$) and α, related to Figure S1F-G**

| Welch's t-test of Log($D_{100ms}$) and α (alpha) | | | | | | | |
|---|---|---|---|---|---|---|---|
| **variable** | **group1** | **group2** | **mean_group1** | **mean_group2** | **t_statistic** | **p_value** | **significant** |
| **log_D** | Uninfected (100ms) | HSV-1 (100ms) | -0.919241291 | -0.296410801 | -90.6221729 | p<0.0001 | TRUE |
| **alpha** | Uninfected (100ms) | HSV-1 (100ms) | 0.589630083 | -0.296410801 | -25.8801123 | p<0.0001 | TRUE |
| **log_D** | Uninfected (200ms) | HSV-1 (200ms) | -1.541429277 | -0.940090921 | -61.59177012 | p<0.0001 | TRUE |
| **alpha** | Uninfected (200ms) | HSV-1 (200ms) | 0.736409203 | 0.75195894 | -4.329269524 | p<0.0001 | TRUE |

**Table 5: Welch's t-test of Log($D_{100ms}$) and α, related to Figure S3H-I**

| **Welch's T-test of Log($D_{100ms}$) and α (alpha)** | | | | | | | |
|---|---|---|---|---|---|---|---|
| **variable** | **group1** | **group2** | **mean_group1** | **mean_group2** | **t_statistic** | **p_value** | **significant** |
| **log_D** | ICP4_none | ICP4_dox | -0.507744888 | -0.345600664 | -22.13672406 | <0.0001 | TRUE |
| **aexp** | ICP4_none | ICP4_dox | 0.672656438 | 0.69236511 | -7.197036553 | <0.0001 | TRUE |
| **log_D** | ICP8_none | ICP8_dox | -0.283165239 | -0.244901402 | -4.492237037 | <0.0001 | TRUE |
| **aexp** | ICP8_none | ICP8_dox | 0.625674373 | 0.602967593 | 6.590069373 | <0.0001 | TRUE |

**Table 6: Welch's t-test of Log($D_{100ms}$) and α, related to Figure S5C-E**

| Welch's T-test of Log(D) and α (alpha) | | | | | | | |
|---|---|---|---|---|---|---|---|
| **variable** | **group1** | **group2** | **Mean group1** | **Mean group2** | **t_statistic** | **p_value** | **significant** |
| **log_D** | ICP4_dox_40Hz | ICP4_dox_4Hz | -1.64538787 | -4.427938942 | 179.046472 | p<0.0001 | TRUE |
| **aexp** | ICP4_dox_40Hz | ICP4_dox_4Hz | 0.4284579 | 0.405969499 | 5.27418607 | p<0.0001 | TRUE |
| **log_D** | ICP4_dox_4Hz | ICP4_un_4Hz | -4.427938942 | -4.705282091 | 12.6294074 | p<0.0001 | TRUE |
| **aexp** | ICP4_dox_4Hz | ICP4_un_4Hz | 0.405969499 | 0.366568018 | 6.59304752 | p<0.0001 | TRUE |
| **log_D** | ICP4_dox_40Hz | ICP4_un_40Hz | -1.64538787 | -1.81912593 | 24.4329352 | p<0.0001 | TRUE |
| **aexp** | ICP4_dox_40Hz | ICP4_un_40Hz | 0.4284579 | 0.378759876 | 22.5996659 | p<0.0001 | TRUE |

**Table 7: One-way ANOVA for Log($D_{100ms}$) and α, related to Figure S6D-E**

| | | | | | | |
|---|---|---|---|---|---|---|
| **One-way ANOVA of Log($D_{100ms}$)** | | | | | | |
| **Source** | **Sum of squares** | **Degrees of freedom** | **F-statistic** | **PR(>F)** | | |
| **Between Groups** | 20844.7343 | 5 | 4703.52299 | <0.0001 | | |
| **Within Groups** | 347675.262 | 392257 | | | | |
| **Multiple Comparison of Means Log($D_{100ms}$) — Tukey HSD, FWER=0.02** | | | | | | |
| **Group 1** | **Group 2** | **MeanDiff** | **p-adj** | **lower** | **upper** | **reject** |
| HSV-1 | Uninfected | -0.3986 | <0.0001 | -0.4148 | -0.3824 | True |
| HSV-1 | HSV-1 (confluent) | -0.0718 | <0.0001 | -0.0861 | -0.0574 | True |
| HSV-1 | HSV-1 + sorbitol | -0.333 | <0.0001 | -0.3484 | -0.3175 | True |
| **One-way ANOVA of α (alpha)** | | | | | | |
| **Source** | **Sum of squares** | **Degrees of freedom** | **F-statistic** | **PR(>F)** | | |
| **Between Groups** | 626.433001 | 5 | 968.651285 | <0.0001 | | |
| **Within Groups** | 50735.0237 | 392257 | | | | |
| **Multiple Comparison of Means α (alpha) — Tukey HSD, FWER=0.02** | | | | | | |
| **Group 1** | **Group 2** | **MeanDiff** | **p-adj** | **lower** | **upper** | **reject** |
| HSV-1 | Uninfected | -0.0896 | <0.0001 | -0.0958 | -0.0834 | True |
| HSV-1 | HSV-1 (confluent) | 0.0038 | 0.2198 | -0.0017 | 0.0093 | False |
| HSV-1 | HSV-1 + sorbitol | -0.036 | <0.0001 | -0.0419 | -0.0301 | True |

**Table 8: Independent t-test of individual total condensate intensity, related to Figure S7B**

| Independent t-test of individual total condensate intensity | | | | | | | |
|---|---|---|---|---|---|---|---|
| **Group 1** | **Group 2** | **t/t0** | **Group 1 Mean** | **Group 2 Mean** | **t-statistic** | **p-value** | **Reject** |
| ICP4 + Dox | ICP4 Untreated | 0 | 1.819160393 | 0.753261776 | 3.163439 | 0.001932 | TRUE |
| ICP4 + Dox | ICP4 Untreated | 1 | 1 | 1 | - | - | - |
| ICP4 + Dox | ICP4 Untreated | 2 | 4.81805612 | 7.106141591 | -1.347765 | 0.179021 | FALSE |
| ICP4 + Dox | ICP4 Untreated | 3 | 10.0273899 | 9.503824831 | 0.281311 | 0.778672 | FALSE |
| ICP4 + Dox | ICP4 Untreated | 4 | 16.61818744 | 12.52824829 | 1.934216 | 0.053994 | FALSE |
| ICP4 + Dox | ICP4 Untreated | 5 | 21.52416715 | 15.09167394 | 2.480298 | 0.013659 | TRUE |
| ICP4 + Dox | ICP4 Untreated | 6 | 23.18740802 | 17.59375055 | 1.968003 | 0.049969 | TRUE |
| ICP4 + Dox | ICP4 Untreated | 7 | 26.33164749 | 17.78268277 | 2.942848 | 0.003497 | TRUE |
| ICP4 + Dox | ICP4 Untreated | 8 | 27.83867205 | 18.88759448 | 2.852948 | 0.004631 | TRUE |
| ICP4 + Dox | ICP4 Untreated | 9 | 28.87550429 | 18.94321931 | 3.269722 | 0.001199 | TRUE |
| ICP4 + Dox | ICP4 Untreated | 10 | 29.31237069 | 19.93860843 | 2.846943 | 0.004718 | TRUE |
| ICP4 + Dox | ICP4 Untreated | 11 | 28.80452435 | 20.68667919 | 2.368969 | 0.018507 | TRUE |
| ICP4 + Dox | ICP4 Untreated | 12 | 30.07036799 | 20.7571277 | 2.774789 | 0.005872 | TRUE |
| ICP4 + Dox | ICP4 Untreated | 13 | 33.00525778 | 22.02540707 | 2.944087 | 0.003495 | TRUE |
| ICP4 + Dox | ICP4 Untreated | 14 | 33.40268396 | 22.07812092 | 2.978785 | 0.003136 | TRUE |
| ICP4 + Dox | ICP4 Untreated | 15 | 33.76764025 | 21.75813273 | 3.213572 | 0.001454 | TRUE |
| ICP4 + Dox | ICP4 Untreated | 16 | 33.71064882 | 22.06184513 | 3.114063 | 0.002028 | TRUE |
| ICP4 + Dox | ICP4 Untreated | 17 | 33.00281889 | 22.60488145 | 2.662139 | 0.008188 | TRUE |
| ICP4 + Dox | ICP4 Untreated | 18 | 32.06660885 | 22.55331309 | 2.447612 | 0.014948 | TRUE |
| ICP4 + Dox | ICP4 Untreated | 19 | 31.33033865 | 22.8993569 | 2.200202 | 0.028607 | TRUE |
| ICP4 + Dox | ICP4 Untreated | 20 | 31.64387065 | 21.98773332 | 2.563329 | 0.010864 | TRUE |
| ICP4 + Dox | ICP4 Untreated | 21 | 32.34620488 | 21.48450489 | 2.831936 | 0.004936 | TRUE |
| ICP4 + Dox | ICP4 Untreated | 22 | 32.73364426 | 21.88897094 | 2.901113 | 0.003988 | TRUE |
| ICP4 + Dox | ICP4 Untreated | 23 | 31.96076 | 22.3114988 | 2.60918 | 0.009536 | TRUE |
| ICP4 + Dox | ICP4 Untreated | 24 | 31.8754685 | 23.55497277 | 2.161496 | 0.031504 | TRUE |
| ICP4 + Dox | ICP4 Untreated | 25 | 31.63797667 | 23.19155166 | 2.211753 | 0.027785 | TRUE |
| ICP4 + Dox | ICP4 Untreated | 26 | 31.74983522 | 23.64361409 | 2.056689 | 0.040646 | TRUE |
| ICP4 + Dox | ICP4 Untreated | 27 | 29.98339235 | 22.37634964 | 2.042356 | 0.042082 | TRUE |
| ICP4 + Dox | ICP4 Untreated | 28 | 29.97309399 | 22.98469628 | 1.785149 | 0.075362 | FALSE |
| ICP4 + Dox | ICP4 Untreated | 29 | 29.54011714 | 22.83062444 | 1.748667 | 0.081503 | FALSE |

**Table 9: Welch's t-test of Log($D_{50min}$) and α, related to Figure S7D**

| Welch's T-test of Log($D_{50min}$) and α (alpha) | | | | | | | |
|---|---|---|---|---|---|---|---|
| **variable** | **group1** | **group2** | **mean_group1** | **mean_group2** | **t_statistic** | **p_value** | **significant** |
| **log_D** | Dox | Untreated | -3.250499965 | -3.402532609 | 5.248803527 | p<0.0001 | TRUE |
| **aexp** | Dox | Untreated | 0.892315691 | 0.876228314 | 0.826788569 | 0.40849 | FALSE |
